# Gotree/Goalign : Toolkit and Go API to facilitate the development of phylogenetic workflows

**DOI:** 10.1101/2021.06.09.447704

**Authors:** Frédéric Lemoine, Olivier Gascuel

## Abstract

Besides computer intensive steps, phylogenetic analysis workflows are usually composed of many small, reccuring, but important data manipulations steps. Among these, we can find file reformatting, sequence renaming, tree re-rooting, tree comparison, bootstrap support computation, etc. These are often performed by custom scripts or by several heterogeneous tools, which may be error prone, uneasy to maintain and produce results that are challenging to reproduce. For all these reasons, the development and reuse of phylogenetic workflows is often a complex task. We identified many operations that are part of most phylogenetic analyses, and implemented them in a toolkit called Gotree/Goalign. The Gotree/Goalign toolkit implements more than 120 user-friendly commands and an API dedicated to multiple sequence alignment and phylogenetic tree manipulations. It is developed in Go, which makes executables efficient, easily installable, integrable in workflow environments, and parallelizable when possible. This toolkit is freely available on most platforms (Linux, MacOS and Windows) and most architectures (amd64, i386). Sources and binaries are available on GitHub at https://github.com/evolbioinfo/gotree, Bioconda, and DockerHub.

## INTRODUCTION

Increase in computer power and development of bioinformatics methods that handle very large datasets make it possible to perform phylogenetic analyses at an unprecedented scale. For example, it is common to perform phylogenetic studies involving several thousand trees or several thousand taxa (*e*.*g*. (1) or (2)). Such studies often present and run complex pipelines involving many steps and several tools and scripts, which makes them hard to describe and share. This is especially true with current Covid-19 pandemics and the diverse SARS-CoV-2 data analysis workflows that flourish, which need to manipulate sequences, alignments, and phylogenetic trees in an easy and automated way (*e*.*g*. https://github.com/roblanf/sarscov2phylo/).

Figure 1 displays an example of a phylogenetic workflow inspired from (3) and described in details in the Results section. The main objective of this workflow is to analyse a phylogenomic dataset to infer a species tree and compare it to a reference tree. It contains 19 steps among which the majority is not constituted by usual computer intensive tasks (e.g. tree inference and multiple sequence alignment), but rather by alignment and tree manipulations such as downloading, renaming, reformating, comparing, rerooting, annotating, etc. These tasks are (i) repetitive (found in many similar workflows), (ii) tedious to implement (many ways to do it), and (iii) error prone, which makes this workflow difficult to implement, describe and reproduce.

**Figure 1.**
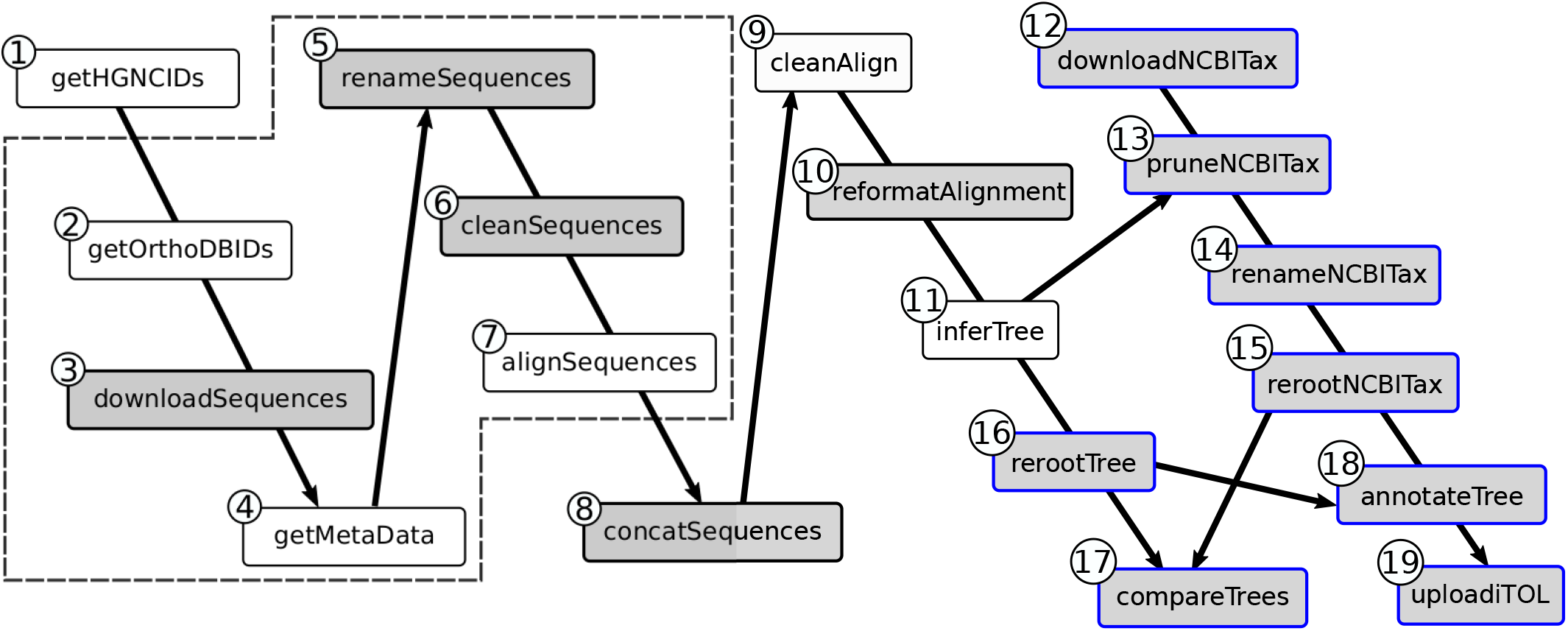
Structure of our use case phylogenomic workflow. It is made of 19 steps (processes), represented as boxes in the figure. Gray boxes represent processes performed using Goalign, and Gray boxes with blue contour represent processes performed using Gotree. Processes are linked by lines if results of the upstream process are needed for the execution of the downstream process. The steps represented by boxes surrounded by the dashed line are executed for each gene in parallel. The steps are the following: **1)** Match RefSeq, NCBI and HGNC (4) gene identifiers; **2)** Get OrthoDB (5) identifiers of orthologous groups corresponding to HGNC identifiers; **3)** Download sequences of each group; **4)** Get species name of each sequence (from OrthoDB id); **5)** Rename the sequences using the species names; **6)** Clean the sequences (*e*.*g*., removing special characters); **7)** Align the sequences (MAFFT (6)); **8)** Concatenate all alignments in a single large alignment; **9**) Clean the alignment (BMGE (7)); **10)** Reformat the alignment into Phylip format; **11)** Infer the phylogenetic tree (IQ-TREE (8)); **12)** Download the NCBI taxonomy in Newick format; **13)** Keep only the 25 species of interest from NCBI taxonomy; **14)** Change species names that differ between OrthoDB and NCBI taxonomy; **15-16)** Reroot NCBI taxonomy and inferred tree; **17)** Compare both trees in terms of common bi-partitions; **18)** Annotate inferred tree with NCBI taxonomy clades; **19)** Upload the annotated tree to iTOL (9).

That is why we think a common framework was needed to help implement such analyzes in a concise and reproducible way.

Several tools, either in the form of command line or APIs, already exist to manipulate phylogenetic trees and/or alignments. For example, Newick-utilities (10) is a command line tool dedicated to phylogenetic tree manipulation in newick format. ETE toolkit (11) is a Python API for the analysis and visualization of trees. Ape (12) is a package dedicated to the analysis of phylogenetics and evolution in the R language. Buddy Suite (13) is another python framework (command line) to manipulate phylogenetic trees and alignments. BIO++ (14) is a C++ library for the analysis of sequences and trees. Lastly, Phyx (15) is a command line tool written in C++ that performs phylogenetic analyses on trees and sequences. Some of these tools (*e*.*g*. ape, ETE toolkit, BIO++) are mainly developer oriented and may be difficult to use for non-programmer phylogenetic analysts who are not familiar with R, python or C++. Others are command line only (*e*.*g*. Newick-utilities), and may be of limited use for developers who want to use part of them in their own programs. They are all implemented either in C, C++, Python or R. To our knowledge, no phylogenetic tree toolkit currently exists, which mix alignment and tree manipulation, in a command line environment and via an API, and proposes a large diversity of commands. In particular, none exists for the Go programming language, which is known to be simple and efficient, and is rising in bioinformatics as demonstrated by the development of several bioinformatics libraries and tools in Go such as biogo (16), Vcfanno (17) or Singularity (18).

In this context, we developed Gotree/Goalign, a toolkit dedicated to burdensome and repetitive phylogenetic tasks, which (i) consists of two user-friendly executables, `gotree` and `goalign`, integrating state of the art phylogenetic commands and requiring no programming skills to use, (ii) provides a set of chainable commands that are integrable in workflows (*e*.*g*. Nextflow (19) or Snakemake (20)), (iii) is straightforward to install via static binaries available for most platforms and architectures, and (iv) provides a public API accessible to developers wanting to manipulate phylogenetic trees and multiple alignments in Go.

The Gotree/Goalign toolkit is already used in several public workflows such as Grapevine (the phylogenetic pipeline for the COG-UK project https://github.com/COG-UK/grapevine), ARTIC-EBOV (The ARTIC Ebola virus phylogenetic analysis protocol https://github.com/artic-network/artic-ebov), KOVID-TREES-NF (a Nextflow (19) workflow for SARS-CoV-2 phylogenetics https://github.com/MDU-PHL/kovid-trees-nf), bioconvert (a generic bioinformatic file format converter https://github.com/bioconvert/bioconvert), and in phylogenetic studies (21, 22, 23, 24, 25, 26, 27).

## MATERIAL AND METHODS

### Alignment manipulation

Goalign implements more than 60 commands to manipulate sequences and multiple sequence alignments. A subset of these commands is given in Table 1.

**Table 1.** Subset of commands implemented in Gotree/Goalign.

| Goalign Command | Description |
| --- | --- |
| reformat | Converts between Nexus, Clustal, Fasta and Phylip |
| rename | Modifies or cleans sequence names |
| clean | Removes sites/sequences |
| codonalign | Aligns by codons using a protein alignment |
| concat | Concatenates several alignments |
| mask | Masks parts of the alignment |
| translate | Translates input alignment in amino-acids |
| extract | Extracts subsequences from input alignments |
| build seqboot | Builds bootstrap alignments |
| compute distance | Computes distance matrix |
| stats | Computes statistics about input alignments |
| Gotree Command | Description |
| reformat | Converts between Newick, Nexus and PhyloXML |
| prune | Removes user-defined tips |
| collapse | Collapses branches |
| reroot | Reroots trees |
| compare trees | Compares tree topologies (bipartitions) |
| compute support | Computes branch supports (31) and (32) |
| stats | Computes statistics about input trees |
| matrix | Computes patristic distance matrix |
| upload itol | Uploads a tree to iTOL (9) |

The most obvious (and maybe the most widely used) commands reformat an input sequence file into any supported format, which is essential to integrate different tools in the same analysis workflow. The supported input and output formats are: Fasta, Phylip, Clustal, Nexus, and tool specific formats (e.g. TNT and PaML input formats). It is worth noting that Goalign supports compressed input files (.gz, .bz, and .xz) and remote files through http(s).

A second kind of commands is designed to extract summary statistics about input alignments, such as the length, the number of sequences, the frequency of each character, the list of sequence names, etc.

The third kind of commands aims at modifying input alignments (e.g. extract sequences or sites, clean, translate, rename, shuffle, mask, concatenate, append, etc.) or producing new alignments (build bootstrap alignment, align by codon, align pairwise, etc.)

Finally, some commands are designed to compute distance matrices for nucleotidic and proteic alignments. To this aim, Goalign implements main evolutionary nucleotidic evolutionary models (JC, K2P, F81, F84 and TN93) and amino-acid matrices (DAYHOFF, JTT, MtRev, LG, WAG). Distances between amino-acid sequences are computed using maximum likelihood as in PhyML(28) and FastME(29).

### Tree manipulation

Gotree implements more than 60 commands dedicated to the manipulation of phylogenetic trees (see Table 1 for a list of a few of them).

As for Goalign, Gotree reformatting commands are the first to be used in phylogenetic workflows and constitute the glue between all phylogenetic tools, which often support heterogeneous, specific formats. Gotree supports the following input and output formats: Newick, Nexus and PhyloXML. Moreover, Gotree supports compressed input files (.gz), and remote files from any urls or from dedicated servers (iTOL (9) and Treebase (30), See Supp. Text 1 for examples of such commands).

The second kind of commands implemented in Gotree produces summary statistics about input phylogenetic trees, such as the number of tips, number of branches, average branch length and branch support, sum of branch lengths, number of cherries, tree balance indices (Colless, Sackin), and patristic distance matrix.

A third kind of commands modifies the input tree, e.g., modifying branch lengths and supports, collapsing branches by length or support, removing lists of tips, rerooting the tree in different ways, etc.

Other commands generate new trees or new data on the trees, such as bootstrap support computations (FBP (31) and TBE (32)), random tree generation using different models, etc. The last kind of commands compares trees, for example comparing topologies (bipartition distance), or comparing tip names, etc.

Put together, these commands provide the user with a large panel of possibilities adapted to many different situations, a few of which are described in the Results section.

### Implementation of Gotree/Goalign

The Gotree/Goalign toolkit is implemented in the Go (https://golang.org/) programming language, and thus takes advantage of (i) being available on major operating systems (Windows, Linux, MacOS), (ii) the richness of the Go standard library and the massive amount of available packages

(http, hashmaps, compression, regex, channels, concurrency, gonum, etc.), (iii) the distributed nature of Go packages, providing phylogenetic packages easily accessible to any developers.

It consists of two executables `gotree` and `goalign` providing all the commands in the same spirit as `git` or `docker`.

All the commands are implemented in the Unix mindset, having one atomic command per task, reading data from the standard input and writing results on the standard output when possible.

### Integration in workflows

The Gotree/Goalign toolkit has been developed to be easily integrated in phylogenetic workflows, using main workflow managers such as Nextflow(19) or Snakemake(20). Two main characteristics facilitate this behavior.

First of all, Gotree/Goalign consists of a large set of command line tools that can be easily chained, and that avoid manual interventions as much as possible. This feature makes them easily usable in workflows as all commands are atomic, easily citable, describable and reproducible.

Moreover, they are made available through three channels that are compatible with the diversity of use in workflow managers, *i*.*e*., multi-plateform executables, Docker and Singularity containers, and Bioconda packages.

## RESULTS - USE CASE IN PHYLOGENOMICS

In this use case, we analyze a phylogenomic dataset inspired from (3), in which the authors analyze a set of 1,730 genes from primates. They infer the species tree either from individual gene trees using ASTRAL III (33) or from gene concatenation using maximum likelihood. Our use case is inspired from the concatenation study, inferring a species tree from available groups of primate orthologous proteins in OrthoDB (5) after having mapped the RefSeq identifiers of the 1,730 analyzed genes to their OrthoDB counterpart.

The workflow is displayed in more detail in Fig. 1 (code available at https://github.com/evolbioinfo/gotree usecase and in Supp. Data 1), and consists of the following main steps (details of the tools and options are given in Supp. Fig. 1): (i) Mapping RefSeq, HGNC (4) and OrthoDB identifiers ; (ii) Retrieving each group of orthologous proteins in OrthoDB if they are present in at least 90% of the 25 primate species (Supp. Data 2) and in a single copy (this results in 1,315 orthologous groups, listed in Supp. Data 3); (iii) For each orthologous group, downloading the sequences of all proteins and their associated metadata; (iv) Renaming, cleaning and aligning sequences using MAFFT (6); (v) Concatenating the alignments of the 1,315 orthologous groups (this results in a single alignment of 674,089 amino-acids); (vi) Cleaning the alignment using BMGE (7) (this keeps 516,999 sites); (vii) Inferring the species tree using IQ-TREE (8); (viii) Downloading the NCBI taxonomy; (ix) Comparing the inferred tree with the NCBI taxonomy; and (x) Uploading both trees to iTOL (9). Each step and their parameters are described in detail in Supp. Fig. 1.

Since it involves many very different steps, this example is challenging to implement and to run fully automatically, without manual intervention. Apart from tree inference and multiple sequence alignment, most of the operations required to develop this workflow are available in Gotree/Goalign toolkit and are straightforward to execute (Supp. Fig. 1). The implementation of the workflow, made of 19 steps and *∼* 350 lines of code, is fully homogeneous in terms of file formats and input/outputs. Furthermore, the ability to run the full analysis, from the input data to the output tree visualization, without manual intervention facilitates its description and reproducibility.

The resulting primate phylogeny, given in Fig. 2, is in almost complete agreement with the NCBI taxonomy, with 15 out of 16 branches of the NCBI taxonomy found in our tree. It is worth noting that the only conflicting branch (in red), grouping Callitrix and Aotus, has a lower bootstrap support than other branches and is also present in the tree inferred using maximum likelihood from gene concatenation in (3). In fact, our topology and theirs (3) are identical.

**Figure 2.**
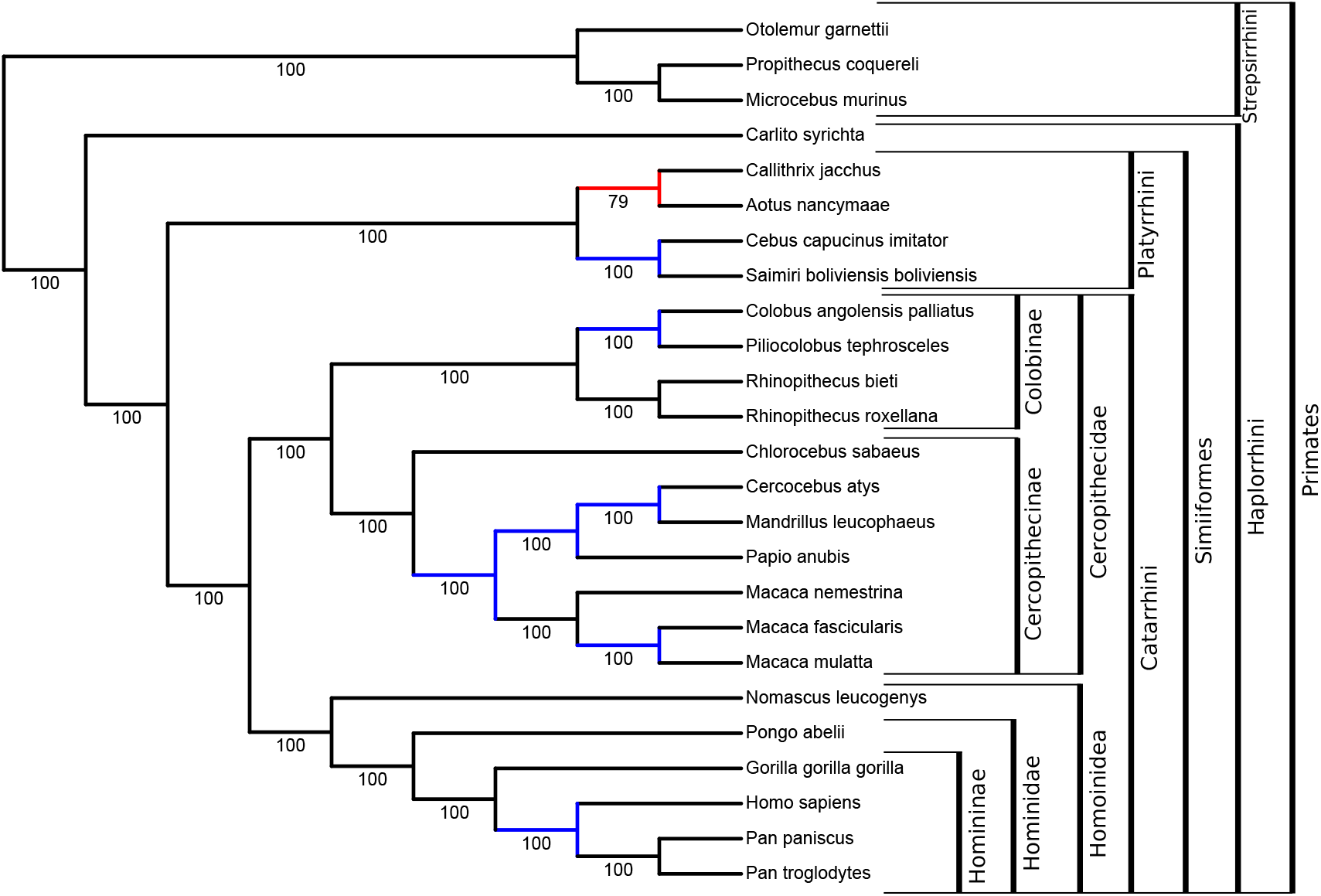
Tree inferred from the concatenation of 1,315 orthologous proteins in 25 primates using maximum likelihood. Visualisation of tree topology with branch supports comes from iTOL (9) after upload by Gotree. Branch colors and clade annotations have been added independently. The red branch is the only one that contradicts the NCBI taxonomy, and blue branches are resolved in the inferred tree, but unresolved in the NCBI taxonomy. The topology is identical to the tree inferred with maximum likelihood from gene concatenation in (3).

## DISCUSSION

We developed the Gotree/Goalign toolkit to simplify the manipulation of phylogenetic trees and alignments, and to facilitate the development of reproducible phylogenetic workflows. Importantly, it is not a wrapper around major tree inference and multiple sequence alignment software, *i*.*e*., it does not take care of aligning sequences and inferring trees, but rather takes care of all the other complex, tedious and numerous tree and sequence operations that are necessary in phylogenetic workflows. The Gotree/Goalign toolkit is developed in Go and is easily installable on major operating systems and provides a public API usable in any Go project.

It is already used in several projects, and we are confident that Gotree/Goalign will be able to build a commmunity interested in adding new functionnalities.

## ACKNOWLEDGEMENTS

First, we thank the Gotree/Goalign users for their feedbacks, which allow us to add new functionnalities and improve existing ones. We also thank all the *Evolutionary Bioinformatics* lab: Anna Zhukova, Marie Morel, Luc Blassel, and Jakub Voznica for testing and using Gotree/Goalign extensively.

