## Supplementary Information for "Gotree/Goalign : Toolkit and Go API to facilitate the development of phylogenetic workflows"

---

---

**Supp. Text 1: Examples of Gotree/Goalign commands** **pp. 2-4**

**Supp Figure 1: Representation of the use case workflow and command templates** **pp. 5-6**

**Supp. Data 1: Nextflow implementation of the use case** **pp. 7-8**

**Supp. Data 2: List of analyzed primate species** **pp. 9**

**Supp. Data 3: List of 1,315 orthologous groups from OrthoDB** **pp. 10-15**

### Supplementary Text 1: Examples of Gotree/Goalign commands

The comprehensive list of Gotree/Goalign commands is given on their respective GitHub repositories:
https://github.com/evolbioinfo/gotree/blob/master/docs/index.md
https://github.com/evolbioinfo/goalign/blob/master/docs/index.md

1) Reformatting a tree from newick to nexus[1]

```
gotree reformat nexus -i itol://129215302173073111930481660
```

The input tree is directly downloaded from iTOL, using its identifier and reformatted in Newick locally.

2) Reformatting an alignment from Fasta to Phylip[1]

```
goalign reformat phylip -i https://github.com/evolbioinfo/goalign/raw/master/tests/data/test_xz.xz
```

The input alignment is automatically downloaded from a remote server, and locally reformatted to Phylip.

3) Display basic summary statistics of a tree from TreeBase[2] :

```
gotree stats --format nexus -i treebase://Tr61953
```

The input tree is directly downloaded from TreeBase, and the following summary statistics are displayed: the number of nodes, tips and edges, the average and total branch length, the average and median support, the number of cherries, and the Colless and Sackin tree balance indices (if rooted).

4) Computing basic summary statistics of a tree from iTOL[2]

```
gotree stats -i itol://129215302173073111930481660
```

This performs the same operation as the previous command, but after downloading the input tree from iTOL.

5) Computing basic summary statistics on a remote alignment[3]

```
goalign stats -i https://github.com/evolbioinfo/goalign/raw/master/tests/data/test_xz.xz
```

The input alignment is downloaded from a remote server, and the following summary statistics are displayed: the length of the alignment, the number of sequences, the average number of different characters per site, the number of variable sites, and the number of occurrences and the frequency of each character (nucleotide or amino-acid).

6) Drawing a tree in the console[4]

```
gotree draw text -w 100 -i
   https://github.com/evolbioinfo/gotree/raw/master/tests/data/rand_tree.nw.gz
```

The input tree is downloaded from a remote server, and the tree is displayed in the console in Phylip like text mode. For example:

```
+ Tip4
|
|---- Tip0
|
|                 +--------- Tip5
|              +--|
|              |  |+--------------------- Tip6
|              |  +|
|              |   |       +------ Tip9
|           +--|   +-------|
|           |  |           +--------------------- Tip3
|           |  |
|           |  |   +- Tip7
+-----------|  +---|
            |      |                        + Tip8
            |      +------------------------|
            |                               +-- Tip2
            |
            +---------------------- Tip1
```

[1] https://github.com/evolbioinfo/gotree/blob/master/docs/commands/reformat.md
[2] https://github.com/evolbioinfo/gotree/blob/master/docs/commands/stats.md
[3] https://github.com/evolbioinfo/goalign/blob/master/docs/commands/stats.md
[4] https://github.com/evolbioinfo/goalign/blob/master/docs/commands/draw.md

7) Rerooting a tree[5]

```
gotree reroot outgroup -i
   https://github.com/evolbioinfo/gotree/raw/master/tests/data/rand_tree.nw.gz Tip484 Tip410 Tip36
```

The input tree is downloaded from a remote server, and is rerooted using the given outgroup defined by a set of tips.

8) Collapsing short branches from a tree[6]

```
gotree collapse length -l 0.01 -i
   https://github.com/evolbioinfo/gotree/raw/master/tests/data/rand_tree.nw.gz
```

The tree is downloaded from a remote server, and branches that are shorter than 0.01 are collapsed, producing polytomies.

9) Compute patristic distance matrix[7]

```
gotree matrix -i https://github.com/evolbioinfo/gotree/raw/master/tests/data/rand_tree.nw.gz
```

The tree is downloaded from a remote server, and the patristic distance matrix is computed (summing over the branch lengths along the paths between all pairs of tips).

10) Mask sites from an alignment[8]

```
goalign mask -s 3 -l 10 -i https://github.com/evolbioinfo/goalign/raw/master/tests/data/test_xz.xz
```

The alignment is downloaded from a remote server, and 10 sites from the $4^{th}$ one (indices start at 0) are masked (replaced by Ns or Xs).

11) Filter out sequences from an alignment[9]

```
goalign subset -r -i https://github.com/evolbioinfo/goalign/raw/master/tests/data/test_xz.xz Seq0002
   Seq0003
```

The input alignment is downloaded from a remote server, and all sequences are removed except the ones given in the command line.

12) Filter out sites from an alignment[10]

```
goalign subsites -r -i https://github.com/evolbioinfo/goalign/raw/master/tests/data/test_xz.xz 1 2 3
```

The input alignment is downloaded from a remote server, and all sites are removed except the ones with indices given in the command line.

13) Compute distances between sequences[11]

```
goalign compute distance -i
   https://github.com/evolbioinfo/goalign/raw/master/tests/data/test_distance.phy.gz --phylip -m jc
```

The DNA input alignment is downloaded from a remote server (--phylip is given because the input format is phylip) and the distance matrix is computed using Jukes and Cantor (1969) evolutionary model (it can be pdist, JC, K2P, F81, F84 and TN93 for DNA alignments, and DAYHOFF, JTT, MtRev, LG and WAG for protein alignments).

14) Concatenate several alignments (merging sequences coming from the same species/taxa)[12]

```
goalign concat -i https://github.com/evolbioinfo/goalign/raw/master/tests/data/test_xz.xz
```

[5] https://github.com/evolbioinfo/gotree/blob/master/docs/commands/reroot.md
[6] https://github.com/evolbioinfo/gotree/blob/master/docs/commands/collapse.md
[7] https://github.com/evolbioinfo/gotree/blob/master/docs/commands/matrix.md
[8] https://github.com/evolbioinfo/goalign/blob/master/docs/commands/mask.md
[9] https://github.com/evolbioinfo/goalign/blob/master/docs/commands/subset.md
[10] https://github.com/evolbioinfo/goalign/blob/master/docs/commands/subsites.md
[11] https://github.com/evolbioinfo/goalign/blob/master/docs/commands/compute.md
[12] https://github.com/evolbioinfo/goalign/blob/master/docs/commands/concat.md

```
    -b https://github.com/evolbioinfo/goalign/raw/master/tests/data/test_xz.xz
```

Several alignments are concatenated (from local files or remote servers), *i.e.* sequences from the same taxa are merged into a single sequence. If a sequence is missing in one of the given alignments, it is replaced by gaps.

15) Build bootstrap alignments[13]

```
goalign build seqboot -i https://github.com/evolbioinfo/goalign/raw/master/tests/data/test_xz.xz -n
500 -o boot
```

The input alignment is downloaded from a remote server, and 500 bootstrap replicates are generated locally.

16) Compute bootstrap support[14]

```
gotree compute support fbp -i
https://github.com/evolbioinfo/gotree/raw/master/tests/data/bootstap_inferred_test.nw.gz \
    -b https://github.com/evolbioinfo/gotree/raw/master/tests/data/bootstap_test.nw.gz
```

A reference tree and a set of bootstrap trees are downloaded from a remote server, and bootstrap supports are computed and attached to branches of the reference tree. Both Felsenstein's (FBP) and transfer version (TBE) of the phylogenetic bootstrap are available with: `gotree compute support fbp` and `gotree compute support tbe`).

[13] https://github.com/evolbioinfo/goalign/blob/master/docs/commands/build.md
[14] https://github.com/evolbioinfo/goalign/blob/master/docs/commands/compute.md

#### Supplementary Figure 1: Representation of the use case workflow and command templates

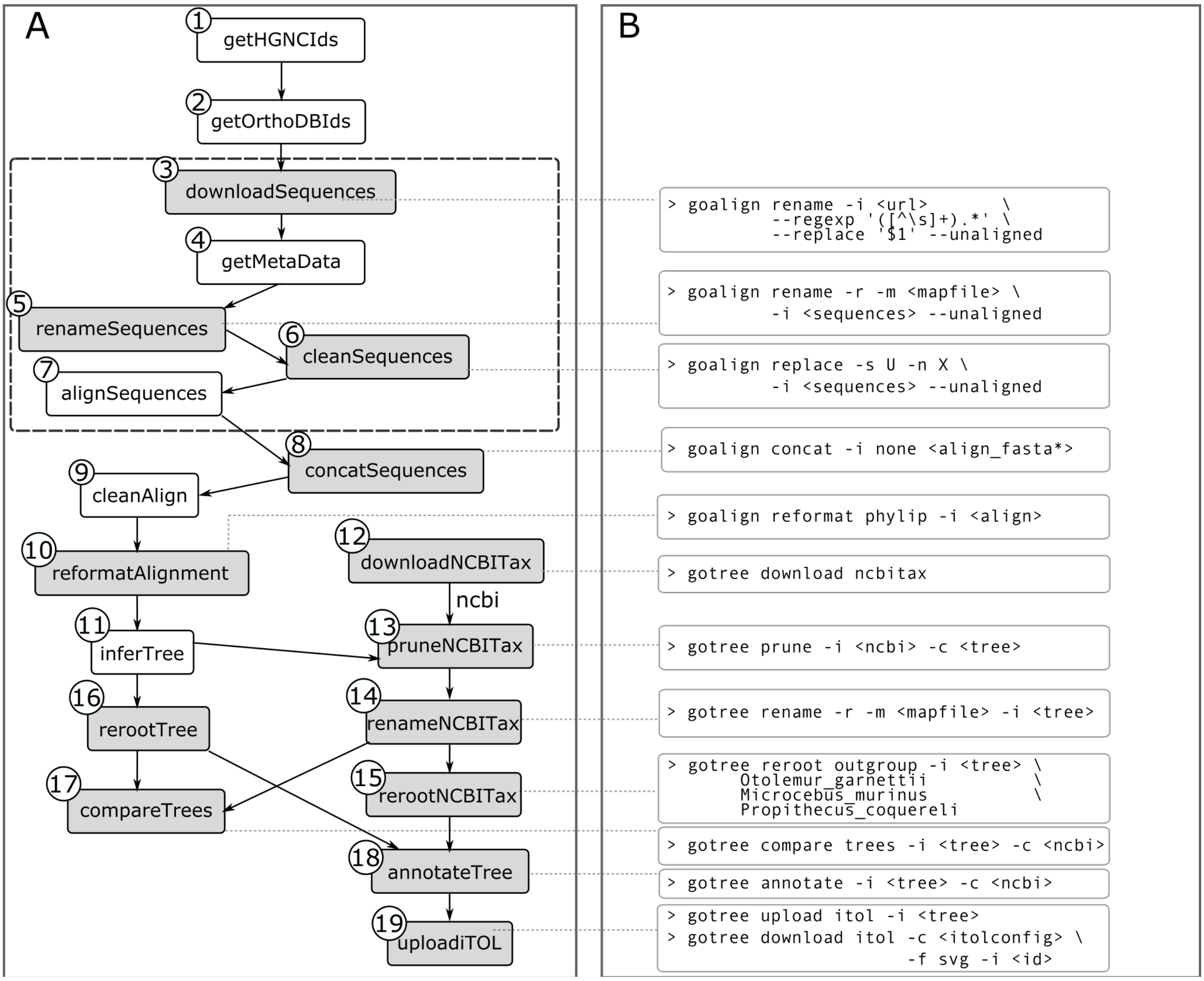


A) This workflow matches RefSeq, HGNC and OrthoDB identifiers from https://doi.org/10.1371/journal.pbio.3000954.s008, and downloads and analyzes 1,315 groups of orthologous proteins having the following characteristics: i) They are shared by at least 90% of the 25 primates (list given below); and ii) only one copy is present in each species (no paralog). The workflow is made of several steps (processes), represented as boxes in the figure. Gray boxes represent steps performed by Gotree/Goalign toolkit. Processes are linked by arrows if results of the upstream process are input of the downstream process. The steps of the workflow, named according to the Nextflow implementation, are the following:

1) **getHGNCIds**: Match RefSeq, NCBI and HGNC (*Eyre et al., NAR 2006*) gene identifiers (dedicated script);
2) **getOrthoDBIds**: Get OrthoDB identifiers of orthologous groups corresponding to HGNC identifiers (orthoDB API);
3) **downloadSequences**: Download sequences of each group and keep only orthoDB ID from the sequence names (Goalign rename command taking the url as input);
4) **getMetaData:** Get species name of each sequence from orthoDB ID (OrthoDB API);
5) **renameSequences**: Rename the sequences using the species names (Goalign);
6) **cleanSequences:** Clean the sequences by removing special characters (Goalign);
7) **alignSequences**: Align the sequences (MAFFT, default options);
8) **concatSequences:** Concatenate all alignments in a single large genomic alignment (Goalign);
9) **cleanAlign:** Clean the alignment (BMGE, with options -t AA -m BLOSUM62 -w 3 -g 0.2 -h 0.5 -b 5);
10) **reformatAlign:** Reformat the alignment into Phylip format (Goalign);
11) **inferTree:** Infer the phylogenetic tree (IQTree, with options -seed 2000 -m MFP -b 100 -nt 20), best model according to Model Finder : JTT+F+R5 (JTT with empirical AA frequencies estimated from the data, and FreeRate model of rates accros sites variability);

12) **DownloadNCBITax:** Download the full NCBI taxonomy in Newick format (Gotree);
13) **pruneNCBITax:** Keep only primate species from NCBI taxonomy (Gotree);
14) **renameNCBITax:** Change species names that differ between orthoDB and NCBI taxonomy (Gotree) ;
15) **rerootNCBITax:** Reroot NCBI taxonomy (Gotree) ;
16) **rerootTree:** Reroot inferred tree using the given outgroup (Gotree);
17) **compareTrees**: Compare both trees in terms of common bi-partitions (Gotree, indicates the number of common and specific bi-partitions);
18) **annotateTree:** Annotate inferred tree with NCBI taxonomy clades (Gotree, indicates for each branch of the tree the closest branch of the NCBI taxonomy in terms of transfer index);
19) **uploadITOL:** Upload the annotated tree to iTOL and store the link to the tree (Gotree).

B) For each step executed using Gotree/Goalign, the template of the command is given in the form "gotree|goalign <command> <subcommand> [options]". Inputs and outputs of most commands / subcommands may be stdin/stdout, allowing for chaining them easily. The first command downloads a set of sequences from OrthoDB and keeps only the OrthoDB identifiers as sequence names; the second command replaces OrthoDB identifiers as sequence names with species names retrieved beforehand.

### Supplementary Data 1: Nextflow implementation of the use case

```
params.nboot = 100
params.seed=2000
params.outpath="results"
params.itolconfig= "data/itol_image_config.txt"
params.refseqid = "data/refseq_ids.txt"
params.ncbirenamefile = "data/ncbi_rename.txt"
params.itolkey="none"
params.itolproject="none"

nboot = params.nboot
seed = params.seed
outpath = file(params.outpath)
itolconfig=file(params.itolconfig)
gene2accession=file("ftp://ftp.ncbi.nlm.nih.gov/gene/DATA/gene2accession.gz")
geneinfo=file("ftp://ftp.ncbi.nlm.nih.gov/gene/DATA/gene_info.gz")
refseqid=file(params.refseqid)
ncbirenamefile=file(params.ncbirenamefile)
itolkey=params.itolkey
itolproject=params.itolproject



// Get the NCBI & HGNC ID Related to the RefSeq protein ID given in
//
https://journals.plos.org/plosbiology/article/file?type=supplementary&id=info:doi/10.1371/journal.pbio.3000954.s008&rev=1
process getNCBIIds{
    publishDir "${outpath}", mode: 'copy'

    input:
    file refseqid
    file gene2accession
    file geneinfo

    output:
    file 'refseq_ids_xref.txt'
    file 'refseq_ids_hgnc.txt' into hgncid

    script:
    """
    add_hgnc.pl $refseqid $gene2accession $geneinfo | sort -u >
refseq_ids_xref.txt
    cut -f 3 refseq_ids_xref.txt | sort -u > refseq_ids_hgnc.txt
    """
}



// Get OrthoDB sequences corresponding to NCBI ID
process getOrthoDBIds{
    maxForks 3

    input:
    val hgnc from hgncid.splitText( by: 1 ).map{ v -> v.trim() }

    output:
    file "ids.txt" into protids,protids2

    script:
    """
    curl
'https://v100.orthodb.org/search?query=${hgnc}&ncbi=0&level=9443&species=9443&singlecopy=1&universal=0.9'|jq '.data | join(",")' | sed 's/"//g' | sed
's/,/\\n/g' > ids.txt
    sleep 1
    """
}

protids2.collectFile(name: "all_orthoid.txt").subscribe{file ->
file.copyTo(outpath.resolve(file.name))}

// Download sequences and metadata
process downloadSequences{
    maxForks 1
    tag "${id}"

    input:
    val id from protids.collectFile(name: 'allids.txt').splitText( by: 1 ).map{
v -> v.trim() }.unique().filter{ it.length() > 0 }

    output:
    tuple val(id), file("sequences.fasta") into sequences

    script:
    """
    goalign rename -i "https://v100.orthodb.org/fasta?id=${id}"  --regexp
'([^\\s]+).*' --replace '\$1' --unaligned > sequences.fasta
    sleep 2
    """
}



// Download meta data
process getMapTable{
    maxForks 1
    tag "${id}"

    input:
    set val(id), file(seq) from sequences

    output:
    set val(id), file(seq), file("map.txt") into mapfile
    file "gene.txt" into genefile

    script:
    """
    wget -O align.tab https://v100.orthodb.org/tab?query=${id}
    cut -f 5,6 align.tab > map.txt
    cut -f 1,2 align.tab | tail -n+2 | sort -u > gene.txt
    """
}
```

```
genefile.collectFile(name: 'genes.txt').subscribe{file ->
file.copyTo(outpath.resolve(file.name))}

// Renaming sequences
process renameSequences{
    tag "${id}"

    input:
    set val(id), file(sequences), file(mapfile) from mapfile

    output:
    file "renamed.fasta" into renamed

    script:
    """
    goalign rename -r -m ${mapfile} -i ${sequences} --unaligned | goalign
rename --regexp " " --replace "_" --unaligned  > renamed.fasta
    """
}



// Cleaning sequences
process cleanSequences{
    input:
    file(sequences) from renamed

    output:
    file "cleaned.fasta" into cleaned

    script:
    """
    goalign replace -s U -n X -i ${sequences} -o cleaned.fasta --unaligned
    """
}



// Aligning sequences
process alignSequences{
    input:
    file cleaned

    output:
    file "aligned.fasta" into alignment

    script:
    """
    mafft --quiet ${cleaned} > aligned.fasta
    """
}



// Concatenating all multiple sequence alignments
process concatSequences {
    input:
    file 'align_fasta' from alignment.collect()

    output:
    file "concat.fasta" into concat

    script:
    """
    goalign concat -o concat.fasta -i none align_fasta*
    """
}

// Cleaning the concatenated multiple sequence alignment (BMGE)
process cleanAlign {
    input:
    file align from concat

    output:
    file "cleanalign.fasta" into cleanalign

    script:
    """
    BMGE -i ${align} -t AA -m BLOSUM62 -w 3 -g 0.2 -h 0.5  -b 5 -of
cleanalign.fasta
    """
}



// Reformatting the alignment in Phylip
process reformatAlignment{
    input:
    file cleanalign

    output:
    file "aligned.phylip" into alignmentphylip

    script:
    """
    goalign reformat phylip -i ${cleanalign} -o aligned.phylip
    """
}

// Inferring the species tree from the concatenated
// multiple sequence alignment
process inferTrueTree{
    publishDir "${outpath}", mode: 'copy'

    input:
    file align from alignmentphylip
    val seed

    output:
    file "tree.nw" into tree, tree2

    script:
    """
```

```
    iqtree -s ${align} -seed ${seed} -m MFP -b 100 -nt ${task.cpus}
    mv *.treefile tree.nw
    """
}

// Rerooting the tree
process rerootTree{
    publishDir "${outpath}", mode: 'copy'

    input:
    file tree from tree2

    output:
    file "rerooted.nw" into reroottree1, reroottree2

    script:
    """
    gotree reroot outgroup -i ${tree} -o rerooted.nw Otolemur_garnettii
Microcebus_murinus Propithecus_coquereli
    """
}

// Downloading the NCBI Taxonomy
process downloadNewickTaxonomy {
    output:
    file "ncbi.nw" into ncbitax

    script:
    """
    gotree download ncbitax -o ncbi.nw
    """
}


// Keep the desired species from the NCBI taxonomy
process pruneNCBITax {

    input:
    file tree from tree
    file map from ncbirenamefile
    file ncbi from ncbitax

    output:
    file "ncbi_pruned.nw" into ncbipruned

    script:
    """
    gotree rename -i ${tree} -m ${map} -r -o tmp
    gotree prune -i ${ncbi} -c tmp -o ncbi_pruned.nw
    """
}

// Rename some tips from the NCBI Taxonomy to match
// orthodb species names
process renameNCBITaxonomy {
    input:
    file ncbi from ncbipruned
    file map from ncbirenamefile

    output:
    file "ncbi_rename.nw" into ncbitaxrename

    script:
    """
    gotree rename -i $ncbi -o ncbi_rename.nw -m $map
    """
}

// Reroot the NCBI taxonomy
process rerootNCBITax {

    input:
    file tree from ncbitaxrename

    output:
    file "ncbi_rerooted.nw" into ncbirerooted1, ncbirerooted2

    script:
    """
    gotree reroot outgroup -i ${tree} -o ncbi_rerooted.nw Otolemur_garnettii
Microcebus_murinus Propithecus_coquereli
    """
}


// Annotating the inferred tree
process annotateTree{
    publishDir "${outpath}", mode: 'copy'

    input:
    file tree from reroottree1
    file ncbi from ncbirerooted1

    output:
    file "annotated.nw" into annotated

    script:
    """
    gotree annotate -i ${tree} -c ${ncbi} -o annotated.nw
    """
}


// Comparing inferred tree and NCBI taxonomy branches
process compareTrees{
    publishDir "${outpath}", mode: 'copy'

    input:
    file tree from reroottree2
    file ncbi from ncbirerooted2
    output:
    file "tree_comparison.txt" into comparison

    script:
    """
    gotree compare trees -i ${tree} -c ${ncbi} > tree_comparison.txt
    """
}

// Uploading tree to ITol &
// Downloading the resulting image
process uploadTree{
    publishDir "${outpath}", mode: 'copy'

    input:
    file tree from annotated
    file itolconfig
    val itolkey
    val itolproject

    output:
    file "tree_url.txt" into iTOLurl
    file "tree_image.svg" into iTOLimage

    script:
    """
    # Upload the tree
    gotree upload itol --name "AnnotatedTree" -i ${tree} --user-id ${itolkey} -
-project ${itolproject} > tree_url.txt
    # We get the iTOL id
    ID=\$(basename \$(cat tree_url.txt ))
    # We Download the image with options defined in data/itol_image_config.txt
    gotree download itol -c ${itolconfig} -f svg -o tree_image.svg -i \$ID
    """
}
```

#### Supplementary Data 2: List of analyzed primate species (OrthoDB ID: 9443)

- Cercocebus atys
- Chlorocebus sabaeus
- Colobus angolensis palliatus
- Macaca fascicularis
- Macaca mulatta
- Macaca nemestrina
- Mandrillus leucophaeus
- Papio anubis
- Piliocolobus tephrosceles
- Rhinopithecus bieti
- Rhinopithecus roxellana
- Gorilla gorilla gorilla
- Homo sapiens
- Pan paniscus
- Pan troglodytes
- Pongo abelii
- Nomascus leucogenys
- Aotus nancymaae
- Callithrix jacchus
- Carlito syrichta
- Cebus capucinus imitator
- Microcebus murinus
- Otolemur garnettii
- Propithecus coquereli
- Saimiri boliviensis boliviensis

### Supplementary Data 3: List of 1,315 orthologous groups from OrthoDB

| | |
|---|---|
| 35432at9443 | Solute carrier family 35 member B1 |
| 36789at9443 | Radial spoke head 1 homolog |
| 27066at9443 | POU domain protein |
| 43408at9443 | Homeobox C12 |
| 32142at9443 | Thioredoxin domain containing 15 |
| 11455at9443 | Microtubule associated protein 6 |
| 12380at9443 | Centrobin, centriole duplication and spindle assembly protein |
| 30512at9443 | Cysteine rich with EGF like domains 2 |
| 23782at9443 | Serpin family A member 11 |
| 29280at9443 | Ganglioside induced differentiation associated protein 1 |
| 39814at9443 | Ras homolog family member J |
| 47676at9443 | Fatty acid binding protein 4 |
| 27275at9443 | dual specificity protein phosphatase 5 |
| 16315at9443 | Dimethylaniline monooxygenase |
| 21933at9443 | INSM transcriptional repressor 2 |
| 33961at9443 | Acyl-CoA binding domain containing 3 |
| 46480at9443 | transmembrane protein 252 |
| 26393at9443 | Arrestin domain containing 2 |
| 48705at9443 | orexigenic neuropeptide QRFP |
| 13878at9443 | selenoprotein O |
| 14869at9443 | NudC domain containing 1 |
| 35926at9443 | Olfactory receptor |
| 13040at9443 | conserved oligomeric Golgi complex subunit 2 isoform X1 |
| 11387at9443 | coiled-coil domain-containing protein 87 |
| 22754at9443 | Gliomedin |
| 25084at9443 | ubiquilin-like protein |
| 48638at9443 | Fatty acid binding protein 1 |
| 36873at9443 | glucagon |
| 36004at9443 | Enoyl-CoA hydratase, short chain 1 |
| 41871at9443 | RNA binding motif protein, X-linked 2 |
| 35629at9443 | Olfactory receptor |
| 35640at9443 | Testis specific serine kinase 3 |
| 44093at9443 | V-set and transmembrane domain containing 2B |
| 22703at9443 | Solute carrier family 19 member 2 |
| 40360at9443 | Chromosome 3 open reading frame 84 |
| 37862at9443 | CD40 ligand |
| 27625at9443 | zinc finger protein 622 |
| 27795at9443 | Family with sequence similarity 187 member B |
| 35153at9443 | POU domain protein |
| 16111at9443 | Phospholipase B-like |
| 27435at9443 | otoconin-90 |
| 18294at9443 | Solute carrier family 32 member 1 |
| 27888at9443 | taste receptor type 2 member 5 |
| 46079at9443 | Chromosome 2 open reading frame 50 |
| 6770at9443 | CTR9 homolog, Paf1/RNA polymerase II complex component |
| 41594at9443 | Ras homolog family member D |
| 24154at9443 | Ladinin-1 |
| 30734at9443 | 5'-nucleotidase, cytosolic IA |
| 32795at9443 | Family with sequence similarity 98 member A |
| 40610at9443 | Homeobox C6 |
| 28498at9443 | Vasopressin V1b receptor |
| 45616at9443 | Amphiregulin |
| 23015at9443 | Aspartate aminotransferase |
| 47568at9443 | glutamyl-tRNA(Gln) amidotransferase subunit C, mitochondrial |
| 39883at9443 | olfactory receptor 8B4 |
| 12729at9443 | DEAQ-box RNA dependent ATPase 1 |
| 46698at9443 | Chromosome 20 open reading frame 85 |
| 23518at9443 | Serpin family C member 1 |
| 43531at9443 | recoverin |
| 47119at9443 | Retinol binding protein 2 |
| 17118at9443 | Kelch like family member 34 |
| 26324at9443 | cathepsin D |
| 30237at9443 | Sodium/potassium-transporting ATPase subunit beta |
| 17598at9443 | Zinc finger protein 662 |
| 6890at9443 | MN1 proto-oncogene, transcriptional regulator |
| 49393at9443 | Secreted LY6/PLAUR domain containing 1 |
| 22638at9443 | Solute carrier family 30 member 1 |
| 48422at9443 | Peptidyl-tRNA hydrolase domain containing 1 |
| 11323at9443 | kelch domain-containing protein 7A |
| 38657at9443 | taste receptor type 2 member 1 |
| 4453at9443 | BUB1 mitotic checkpoint serine/threonine kinase |
| 44903at9443 | BCL2 related protein A1 |
| 16598at9443 | DEF6, guanine nucleotide exchange factor |
| 10037at9443 | Protein kinase C eta type |
| 19881at9443 | Scavenger receptor class B member 2 |
| 28254at9443 | ETS variant 7 |
| 32454at9443 | POU domain protein |
| 39253at9443 | ferritin, mitochondrial |
| 31663at9443 | Phosphatidylinositol 4-kinase type 2 alpha |
| 33818at9443 | transmembrane protein 115 |
| 42070at9443 | Retinol binding protein 1 |
| 39125at9443 | Transmembrane protein 70 |
| 42751at9443 | CD68 molecule |
| 22585at9443 | Bone morphogenetic protein 7 |
| 38043at9443 | protein FAM122A |
| 50804at9443 | Cyclin-dependent kinases regulatory subunit |
| 37220at9443 | Claudin |
| 48796at9443 | Chromosome 15 open reading frame 62 |
| 43761at9443 | Crystallin gamma B |
| 36377at9443 | zinc/RING finger protein 4 |
| 34691at9443 | Gastrulation brain homeobox 2 |
| 19314at9443 | inositol 1,4,5-trisphosphate receptor-interacting protein-like 2 |
| 28692at9443 | Dynein cytoplasmic 2 light intermediate chain 1 |
| 39964at9443 | StAR related lipid transfer domain containing 5 |
| 20753at9443 | PLAG1 like zinc finger 1 |
| 33252at9443 | melanocortin receptor 4 |
| 28453at9443 | Chromosome 2 open reading frame 69 |
| 46047at9443 | Myosin light chain 1 |
| 32564at9443 | UbiA prenyltransferase domain containing 1 |
| 39038at9443 | Yip1 interacting factor homolog A, membrane trafficking protein |
| 17438at9443 | Cholinergic receptor nicotinic delta subunit |
| 49824at9443 | LOW QUALITY PROTEIN: BH3-like motif-containing cell death inducer |
| 35448at9443 | TNF alpha induced protein 6 |
| 42368at9443 | ubiquitin-conjugating enzyme E2 G1 |
| 38397at9443 | Adrenoceptor beta 3 |
| 28601at9443 | Transforming growth factor beta regulator 1 |
| 44885at9443 | Chromosome 8 open reading frame 46 |
| 5241at9443 | Kinesin family member 11 |
| 7082at9443 | tubulin monoglycylase TTLL3 isoform X1 |
| 37714at9443 | Homeobox A10 |
| 15529at9443 | Ribosomal RNA processing 1B |
| 43452at9443 | Protein lin-7 homolog |
| 30848at9443 | Tartrate-resistant acid phosphatase type 5 |
| 41251at9443 | Claudin |
| 14582at9443 | Biotinidase |
| 23630at9443 | forkhead box protein O4 |
| 40694at9443 | alpha-ketoglutarate-dependent dioxygenase alkB homolog 6 isoform X1 |
| 52140at9443 | adipogenesis regulatory factor |
| 17416at9443 | ATPase H+ transporting V1 subunit B1 |
| 39630at9443 | Single-pass membrane protein with coiled-coil domains 1 |
| 38448at9443 | Fc fragment of IgE receptor Ia |
| 37877at9443 | Enoyl-CoA delta isomerase 1 |
| 47021at9443 | Retinol binding protein 5 |
| 27421at9443 | NADH dehydrogenase |
| 18221at9443 | Translocase of outer mitochondrial membrane 70 |
| 26245at9443 | transmembrane protease serine 11D |
| 34770at9443 | RCSD domain containing 1 |
| 48816at9443 | Succinate dehydrogenase complex assembly factor 3 |
| 26703at9443 | Synaptotagmin 13 |
| 45603at9443 | DNL-type zinc finger protein |
| 50242at9443 | splicing factor 3B subunit 5 |
| 31027at9443 | Secreted frizzled related protein 4 |
| 37045at9443 | Junctional adhesion molecule 2 |
| 40827at9443 | B9 domain containing 1 |
| 30646at9443 | Inositol monophosphatase domain containing 1 |
| 40001at9443 | YjeF N-terminal domain containing 3 |
| 21665at9443 | lymphocyte cytosolic protein 2 |
| 19837at9443 | dystrotelin |
| 34299at9443 | G protein-coupled receptor 150 |
| 37113at9443 | N(alpha)-acetyltransferase 30, NatC catalytic subunit |
| 40661at9443 | SS18L1, nBAF chromatin remodeling complex subunit |
| 31118at9443 | MOS proto-oncogene, serine/threonine kinase |
| 29443at9443 | Dol-P-Man |
| 20543at9443 | Matrilin 3 |
| 14740at9443 | Olfactomedin like 2A |
| 39311at9443 | Transmembrane protein 65 |
| 43017at9443 | Ubiquitin conjugating enzyme E2 S |
| 10232at9443 | Phosphoinositide-3-kinase adaptor protein 1 |
| 35841at9443 | Calbindin 2 |
| 26486at9443 | Zinc finger protein 764 |
| 51031at9443 | Chromosome 1 open reading frame 189 |
| 28956at9443 | Ankyrin repeat and sterile alpha motif domain containing 4B |
| 47181at9443 | Cornifelin |
| 28323at9443 | Abhydrolase domain containing 1 |
| 33770at9443 | Potassium channel tetramerization domain containing 13 |
| 43580at9443 | NK6 homeobox 1 |
| 27827at9443 | Cathepsin H |
| 27859at9443 | Major facilitator superfamily domain containing 9 |
| 15419at9443 | Dolichyl-diphosphooligosaccharide--protein glycosyltransferase subunit 1 |
| 37892at9443 | keratin-associated protein 24-1 |
| 34633at9443 | Solute carrier family 35 member G1 |
| 26731at9443 | ameloblastin |
| 45335at9443 | Chromosome X open reading frame 66 |
| 39541at9443 | Fat storage inducing transmembrane protein 1 |
| 8928at9443 | NOP14 nucleolar protein |
| 13279at9443 | Adducin 3 |
| 46086at9443 | Endothelin 2 |
| 34944at9443 | RING finger protein 148 |
| 33502at9443 | Neurogenic differentiation factor |
| 640at9443 | ATP-binding cassette sub-family A member 2 |
| 34488at9443 | Dermokine |
| 28848at9443 | nucleosome assembly protein 1-like 2 |
| 41673at9443 | RAB39B, member RAS oncogene family |
| 22284at9443 | Proteasome 26S subunit, non-ATPase 3 |
| 32631at9443 | secreted frizzled-related protein 2 |
| 37429at9443 | zinc finger CCCH-type antiviral protein 1-like |
| 28764at9443 | Chromosome 1 open reading frame 87 |
| 28700at9443 | Dual specificity protein phosphatase |
| 43545at9443 | Paired like homeobox 2a |
| 46432at9443 | Ubiquitin conjugating enzyme E2 L6 |
| 6836at9443 | mutS protein homolog 4 |
| 28958at9443 | Transmembrane p24 trafficking protein family member 8 |
| 26544at9443 | Tudor domain containing 10 |
| 25495at9443 | AlkB homolog 1, histone H2A dioxygenase |
| 29624at9443 | Replication factor C subunit 5 |
| 30164at9443 | Mesoderm posterior bHLH transcription factor 2 |
| 47093at9443 | Ubiquitously expressed prefoldin like chaperone |
| 40027at9443 | Transmembrane protein 147 |
| 16995at9443 | thrombomodulin |
| 19145at9443 | interferon regulatory factor 2-binding protein 1 |
| 24704at9443 | Nerve growth factor receptor |
| 7006at9443 | Transcriptional regulating factor 1 |
| 12531at9443 | exocyst complex component 8 |
| 796at9443 | Human immunodeficiency virus type I enhancer binding protein 2 |
| 38840at9443 | Kruppel like factor 9 |
| 39801at9443 | Chromosome 16 open reading frame 78 |
| 11160at9443 | Transporter 1, ATP binding cassette subfamily B member |
| 50543at9443 | DnaJ heat shock protein family (Hsp40) member C15 |
| 40757at9443 | sepiapterin reductase |
| 29949at9443 | Solute carrier family 30 member 8 |
| 15117at9443 | WD repeat domain 43 |
| 21072at9443 | Solute carrier family 16 member 2 |
| 46433at9443 | IQ motif containing F2 |
| 38975at9443 | Secretory carrier-associated membrane protein |
| 34362at9443 | Mitochondrial ribosomal protein L2 |
| 38601at9443 | 2-aminoethanethiol dioxygenase |
| 25161at9443 | wnt inhibitory factor 1 |
| 21618at9443 | Lecithin-cholesterol acyltransferase |
| 6127at9443 | Poly |
| 39459at9443 | Stanniocalcin 1 |
| 15357at9443 | Phosphodiesterase 12 |
| 19162at9443 | Ectonucleoside triphosphate diphosphohydrolase 8 |
| 36578at9443 | CutC copper transporter |
| 28847at9443 | Methyltransferase like 14 |
| 46500at9443 | transmembrane protein 160 |
| 30743at9443 | Family with sequence similarity 221 member B |
| 42381at9443 | ciliary neurotrophic factor |
| 17408at9443 | HERV-H LTR-associating protein 1 |
| 48918at9443 | splicing factor 3B subunit 6 |
| 33083at9443 | Olfactory receptor |
| 36066at9443 | V-set and immunoglobulin domain containing 2 |
| 41339at9443 | RAB25, member RAS oncogene family |
| 12429at9443 | Coiled-coil domain containing 142 |
| 45948at9443 | Claudin |

| | |
|---|---|
| 3774at9443 | retinol-binding protein 3 |
| 23820at9443 | LanC like 3 |
| 21424at9443 | WD repeat domain 88 |
| 45285at9443 | PYD and CARD domain containing |
| 24692at9443 | Adrenoceptor beta 2 |
| 38372at9443 | E3 ubiquitin-protein ligase |
| 13333at9443 | SPOC domain containing 1 |
| 42824at9443 | DAN domain family member 5 |
| 19053at9443 | Tyrosine hydroxylase |
| 3739at9443 | golgi glycoprotein 1 |
| 21862at9443 | Lipase |
| 40688at9443 | TP53 regulating kinase |
| 8148at9443 | Spalt like transcription factor 2 |
| 42144at9443 | Stromal cell derived factor 2 like 1 |
| 12721at9443 | Tubulin tyrosine ligase like 12 |
| 37685at9443 | three prime repair exonuclease 2 |
| 5126at9443 | Sodium/potassium-transporting ATPase subunit alpha |
| 31695at9443 | Calcium homeostasis modulator family member 4 |
| 32227at9443 | melanocortin receptor 5 |
| 33178at9443 | Somatomedin B and thrombospondin type 1 domain containing |
| 29720at9443 | 2-oxoglutarate and iron dependent oxygenase domain containing 2 |
| 39406at9443 | NK2 homeobox 6 |
| 48647at9443 | phospholipase A2 |
| 46678at9443 | BTG anti-proliferation factor 2 |
| 28831at9443 | Cytotoxic and regulatory T cell molecule |
| 37098at9443 | Distal-less homeobox 3 |
| 43816at9443 | Mitochondrial ribosomal protein S23 |
| 24320at9443 | uncharacterized protein C3orf30 homolog |
| 47007at9443 | Centrin 3 |
| 40278at9443 | INO80 complex subunit B |
| 34596at9443 | Methylthioribose-1-phosphate isomerase |
| 27237at9443 | actin-like protein 9 |
| 35895at9443 | proline synthase co-transcribed bacterial homolog protein |
| 49672at9443 | EP300-interacting inhibitor of differentiation 2B |
| 38613at9443 | Shisa family member 2 |
| 24600at9443 | Receptor interacting serine/threonine kinase 3 |
| 32125at9443 | F-actin-capping protein subunit alpha-3 |
| 44287at9443 | Lens fiber membrane intrinsic protein |
| 38475at9443 | Glutaredoxin and cysteine rich domain containing 2 |
| 30588at9443 | Opsin 1, short wave sensitive |
| 32731at9443 | Aquaporin 3 (Gill blood group) |
| 15097at9443 | Tripartite motif containing 23 |
| 14840at9443 | CD93 molecule |
| 44435at9443 | B-cell lymphoma/leukemia 10 |
| 32852at9443 | CRK like proto-oncogene, adaptor protein |
| 22012at9443 | Angiopoietin like 3 |
| 15809at9443 | inactive serine/threonine-protein kinase PLK5 |
| 19664at9443 | N-lysine methyltransferase SETD6 |
| 12449at9443 | retrotransposon-derived protein PEG10 isoform 1 |
| 27907at9443 | Annexin |
| 38203at9443 | Vacuolar-sorting protein SNF8 |
| 47824at9443 | mitochondrial import inner membrane translocase subunit Tim22 |
| 40116at9443 | transcription factor MafB |
| 50388at9443 | Interleukin-2 |
| 13044at9443 | Solute carrier family 44 member 3 |
| 12121at9443 | Solute carrier family 39 member 12 |
| 21077at9443 | NADPH oxidase activator 1 isoform X1 |
| 32549at9443 | Ankyrin repeat domain 1 |
| 20322at9443 | Aconitate decarboxylase 1 |
| 35257at9443 | TruB pseudouridine synthase family member 1 |
| 15669at9443 | tetratricopeptide repeat protein 24 |
| 25395at9443 | Mitogen-activated protein kinase |
| 33144at9443 | CTD small phosphatase like |
| 34473at9443 | synapse-associated protein 1 |
| 38717at9443 | CD79a molecule |
| 38350at9443 | Chymase 1 |
| 35415at9443 | Tumor necrosis factor ligand superfamily member |
| 36068at9443 | T cell leukemia homeobox 3 |
| 25396at9443 | GDNF family receptor alpha like |
| 37005at9443 | SLAM family member 7 |
| 44713at9443 | uncharacterized protein C6orf47 homolog |
| 25214at9443 | Microspherule protein 1 |
| 18818at9443 | Matrix metalloproteinase |
| 42464at9443 | Transcription factor SOX |
| 14516at9443 | Growth factor receptor bound protein 10 |
| 39079at9443 | Interleukin 22 receptor subunit alpha 2 |
| 31002at9443 | UTP3, small subunit processome component |
| 18514at9443 | Complement C9 |
| 24030at9443 | Multiple EGF like domains 9 |
| 20604at9443 | NMDA receptor synaptonuclear signaling and neuronal migration factor isoform X1 |
| 30887at9443 | Glycerophosphodiester phosphodiesterase domain containing 3 |
| 38134at9443 | Calcium homeostasis modulator family member 6 |
| 18860at9443 | pre-mRNA-processing factor 19 |
| 12030at9443 | Lipase maturation factor |
| 34348at9443 | Homeobox A2 |
| 42686at9443 | spermatid maturation protein 1 |
| 13287at9443 | sulfhydryl oxidase 2 |
| 27810at9443 | Spectrin repeat containing nuclear envelope family member 4 |
| 20859at9443 | cartilage matrix protein |
| 28751at9443 | NIPA like domain containing 4 |
| 43794at9443 | Clarin 2 |
| 25792at9443 | Nectin cell adhesion molecule 2 |
| 32402at9443 | peroxisome biogenesis factor 13 |
| 28837at9443 | homeobox protein aristaless-like 4 |
| 50261at9443 | LSM4 homolog, U6 small nuclear RNA and mRNA degradation associated |
| 22842at9443 | Cytidine monophosphate N-acetylneuraminic acid synthetase |
| 39073at9443 | Hexamethylene bisacetamide inducible 1 |
| 49344at9443 | Interferon induced transmembrane protein 5 |
| 36727at9443 | TP53 induced glycolysis regulatory phosphatase |
| 25528at9443 | protein FAM81B |
| 20415at9443 | Kremen protein |
| 35468at9443 | zinc finger DHHC-type containing 19 |
| 36806at9443 | Leucine rich repeat containing 58 |
| 46139at9443 | Cytidine deaminase |
| 35986at9443 | ubiquitin-conjugating enzyme E2 Z |
| 34382at9443 | Solute carrier family 51 alpha subunit |
| 21108at9443 | matrix metallopeptidase 8 |
| 35746at9443 | BCDIN3 domain containing RNA methyltransferase |
| 25730at9443 | Choline kinase beta |
| 8722at9443 | Amine oxidase |
| 28772at9443 | TBC1 domain family member 21 |
| 32742at9443 | Dehydrogenase/reductase 7C |
| 30565at9443 | thymidylate synthase |
| 42606at9443 | Non-specific cytotoxic cell receptor protein 1 homolog (zebrafish) |
| 43656at9443 | Claudin |
| 45641at9443 | Chromosome 11 open reading frame 88 |
| 22773at9443 | Ectonucleoside triphosphate diphosphohydrolase 2 |
| 49354at9443 | prefoldin subunit 2 |
| 49128at9443 | Chromosome 11 open reading frame 97 |
| 51627at9443 | uncharacterized protein C11orf71 homolog |
| 36548at9443 | matrilysin |
| 48928at9443 | Thiosulfate sulfurtransferase like domain containing 1 |
| 10158at9443 | Unkempt family zinc finger |
| 11004at9443 | cadherin-15 |
| 46332at9443 | musculin |
| 26001at9443 | cytoskeleton-associated protein 4 |
| 44207at9443 | Protein yippee-like |
| 35441at9443 | ceroid-lipofuscinosis neuronal protein 6 |
| 21383at9443 | interleukin-1 receptor-associated kinase 1 |
| 40715at9443 | Receptor expression-enhancing protein |
| 39236at9443 | Zinc finger protein 414 |
| 30198at9443 | Sulfotransferase |
| 31817at9443 | Pim-2 proto-oncogene, serine/threonine kinase |
| 26160at9443 | beta-1,4-galactosyltransferase 5 |
| 44887at9443 | Proteasome subunit beta type |
| 48472at9443 | Oligodendrocytic myelin paranodal and inner loop protein |
| 50143at9443 | C-C motif chemokine 22 |
| 36234at9443 | Transmembrane protein 150A |
| 29089at9443 | Signal regulatory protein beta 2 |
| 46543at9443 | serine/arginine-rich splicing factor 1 |
| 24851at9443 | Indoleamine 2,3-dioxygenase 2 |
| 9956at9443 | ATP binding cassette subfamily D member 2 |
| 23340at9443 | Coiled-coil domain containing 17 |
| 22819at9443 | reticulon-4 receptor |
| 10211at9443 | ATP binding cassette subfamily B member 7 |
| 40195at9443 | E3 ubiquitin-protein ligase RNF187 |
| 41953at9443 | centromere protein H |
| 48116at9443 | Coiled-coil domain containing 182 |
| 25777at9443 | Phospholipid phosphatase related 2 |
| 44873at9443 | Vacuolar protein sorting 25 homolog |
| 29945at9443 | Translocating chain-associated membrane protein |
| 35117at9443 | Homeobox B13 |
| 42213at9443 | Sugar transporter SWEET |
| 44067at9443 | Adenine phosphoribosyltransferase |
| 17243at9443 | Arylsulfatase B |
| 47700at9443 | refilin B |
| 32073at9443 | ATPase H+/K+ transporting beta subunit |
| 32860at9443 | Metaxin 3 |
| 21943at9443 | Hemopexin |
| 20148at9443 | Leucine rich repeat containing 47 |
| 30676at9443 | Galectin |
| 46599at9443 | Troponin C1, slow skeletal and cardiac type |
| 23857at9443 | Solute carrier family 37 member 4 |
| 45080at9443 | Interleukin-18 |
| 34763at9443 | Ankyrin repeat domain 63 |
| 49061at9443 | H1 histone family member X |
| 45657at9443 | Ribosomal RNA processing 36 |
| 36344at9443 | Phosphomannomutase |
| 42282at9443 | Cilia and flagella associated protein 20 |
| 27286at9443 | IQ motif containing D |
| 30702at9443 | Hyaluronan and proteoglycan link protein 4 |
| 46422at9443 | DnaJ heat shock protein family (Hsp40) member C24 |
| 43745at9443 | Pentaxin |
| 11431at9443 | TLR4 interactor with leucine rich repeats |
| 50851at9443 | Glutamate rich 4 |
| 9618at9443 | Aldehyde dehydrogenase family 16 member A1 |
| 38117at9443 | Phosphoinositide-3-kinase interacting protein 1 |
| 50314at9443 | guanylate cyclase activator 2B |
| 38378at9443 | Thioredoxin related transmembrane protein 1 |
| 8195at9443 | AP-2 complex subunit alpha |
| 39390at9443 | tumor necrosis factor ligand superfamily member 15 |
| 20158at9443 | BEN domain containing 2 |
| 31390at9443 | Free fatty acid receptor 4 |
| 29571at9443 | Testis specific serine kinase 4 |
| 50738at9443 | Serine peptidase inhibitor, Kazal type 7 (putative) |
| 32604at9443 | eukaryotic translation initiation factor 3 subunit G |
| 46200at9443 | Intercellular adhesion molecule 4 (Landsteiner-Wiener blood group) |
| 13733at9443 | Zinc finger protein 599 |
| 24504at9443 | CDC like kinase 2 |
| 39562at9443 | retinoschisin |
| 43935at9443 | Glutathione peroxidase |
| 42305at9443 | transmembrane emp24 domain-containing protein 7 |
| 28018at9443 | Mitochondrial ribosomal protein L37 |
| 24325at9443 | carboxypeptidase B |
| 26427at9443 | Leiomodin 2 |
| 28126at9443 | Mex-3 RNA binding family member A |
| 25044at9443 | kaptin |
| 47970at9443 | Transmembrane protein 207 |
| 14157at9443 | Zinc finger SWIM-type containing 2 |
| 41669at9443 | regulator of G-protein signaling 1 |
| 29328at9443 | Protein Wnt |
| 31535at9443 | G-protein coupled receptor 20 |
| 13317at9443 | Interleukin 27 receptor subunit alpha |
| 27781at9443 | CD5 molecule like |
| 44083at9443 | Sodium voltage-gated channel beta subunit 2 |
| 40137at9443 | Exosome component 5 |
| 6489at9443 | Espin like |
| 51324at9443 | neuropeptide S |
| 725at9443 | lipoxygenase homology domain-containing protein 1 isoform X1 |
| 27737at9443 | Growth differentiation factor 2 |
| 22097at9443 | alpha-(1,3)-fucosyltransferase 4 |
| 18299at9443 | Gap junction protein |
| 44584at9443 | Tetratricopeptide repeat domain 9 |
| 47964at9443 | cystatin-like 1 |
| 51565at9443 | S100 calcium binding protein A12 |
| 49502at9443 | zinc finger protein 593 |
| 41858at9443 | Folliculogenesis specific bHLH transcription factor |
| 20279at9443 | Adenylosuccinate synthetase isozyme 2 |
| 40220at9443 | Homeobox A6 |
| 41211at9443 | Protease associated domain containing 1 |
| 40300at9443 | Quinoid dihydropteridine reductase |
| 34270at9443 | taste receptor type 2 member 39 |
| 19333at9443 | Testis expressed metallothionein like protein |
| 8518at9443 | AE binding protein 1 |
| 33107at9443 | Na(+)/H(+) exchange regulatory cofactor NHE-RF1 |
| 30485at9443 | Arginase |
| 26877at9443 | pre-mRNA-splicing factor RBM22 |
| 28604at9443 | Neuromedin U receptor 1 |

| | |
|---|---|
| 37295at9443 | Solute carrier family 35 member E4 |
| 42439at9443 | Amelotin |
| 46194at9443 | charged multivesicular body protein 4c |
| 33417at9443 | Stomatin like 2 |
| 35347at9443 | Serine protease 57 |
| 27975at9443 | Keratin 20 |
| 22404at9443 | Lactate dehydrogenase D |
| 30562at9443 | nucleosome assembly protein 1-like 3 |
| 51098at9443 | Liver enriched antimicrobial peptide 2 |
| 16490at9443 | EGF like domain multiple 6 |
| 34933at9443 | STIP1 homology and U-box containing protein 1 |
| 6824at9443 | DExD/H-box helicase 58 |
| 33169at9443 | BarH like homeobox 2 |
| 963at9443 | Voltage-dependent N-type calcium channel subunit alpha |
| 48373at9443 | BCL tumor suppressor 7A |
| 33800at9443 | olfactory receptor 13J1 |
| 50950at9443 | Transmembrane protein 35B |
| 31728at9443 | nucleoside diphosphate-linked moiety X motif 19 |
| 9893at9443 | DEAD-box helicase 1 |
| 46447at9443 | charged multivesicular body protein 1b |
| 43195at9443 | C-type lectin domain family 4 member E |
| 25682at9443 | Ankyrin repeat and SOCS box containing 16 |
| 42555at9443 | Fibroblast growth factor |
| 16439at9443 | Major facilitator superfamily domain containing 6 like |
| 31488at9443 | Solute carrier family 10 member 4 |
| 31994at9443 | Solute carrier family 35 member E3 |
| 32897at9443 | Beta-1,3-N-acetylglucosaminyltransferase |
| 36494at9443 | coiled-coil domain containing 28B |
| 34801at9443 | Olfactory receptor |
| 27120at9443 | ST8 alpha-N-acetyl-neuraminide alpha-2,8-sialyltransferase 3 |
| 25362at9443 | transcription factor IIIA |
| 17048at9443 | TBC1 domain family member 24 isoform X1 |
| 23443at9443 | sphingosine-1-phosphate phosphatase 1 |
| 42810at9443 | ADP ribosylation factor like GTPase 14 |
| 50691at9443 | basic leucine zipper transcriptional factor ATF-like |
| 41130at9443 | Fanconi anemia core complex associated protein 24 |
| 37010at9443 | Atonal bHLH transcription factor 1 |
| 33452at9443 | melanocortin receptor 3 |
| 47455at9443 | Profilin |
| 33913at9443 | Ankyrin repeat domain 23 |
| 27619at9443 | G protein-coupled receptor 151 |
| 25333at9443 | One cut domain family member |
| 14362at9443 | Complement C8 beta chain |
| 41732at9443 | Myelin protein zero like 3 |
| 21445at9443 | peptidyl-prolyl cis-trans isomerase FKBP5 |
| 46614at9443 | leucine rich adaptor protein 1-like |
| 29906at9443 | Dual specificity protein phosphatase |
| 13303at9443 | Crooked neck pre-mRNA splicing factor 1 |
| 17868at9443 | uncharacterized protein C2orf81 homolog isoform X1 |
| 40824at9443 | Retina and anterior neural fold homeobox |
| 39909at9443 | Ribosome assembly factor mrt4 |
| 38910at9443 | Homeobox protein |
| 6063at9443 | VPS50, EARP/GARPII complex subunit |
| 39639at9443 | tumor necrosis factor ligand superfamily member 6 |
| 37547at9443 | Tubulin epsilon and delta complex 1 |
| 32910at9443 | VPS26 endosomal protein sorting factor C |
| 44084at9443 | MAGE family member H1 |
| 43559at9443 | Ankyrin repeat domain 22 |
| 39123at9443 | Small nuclear ribonucleoprotein polypeptide A' |
| 42829at9443 | Transmembrane 4 L six family member 20 |
| 27926at9443 | olfactory receptor 1I1 |
| 18378at9443 | T-box 21 |
| 43036at9443 | transmembrane protein 179 |
| 25122at9443 | early growth response protein 1 |
| 18877at9443 | T-complex protein 1 subunit delta |
| 17982at9443 | Aldehyde dehydrogenase 9 family member A1 |
| 23077at9443 | PWP1 homolog, endonuclein |
| 42108at9443 | Ras homolog family member B |
| 40895at9443 | WW domain binding protein 1 |
| 45417at9443 | Palmitoyltransferase |
| 46801at9443 | SCP2 sterol-binding domain-containing protein 1 |
| 30001at9443 | Isthmin 2 |
| 34746at9443 | taste receptor type 2 member 3 |
| 38761at9443 | insulin-like growth factor-binding protein 1 |
| 28815at9443 | Angiotensin II receptor type 2 |
| 46592at9443 | Calmodulin like 3 |
| 28438at9443 | Glucose-6-phosphatase |
| 42421at9443 | MAGE family member F1 |
| 51738at9443 | Chromosome 10 open reading frame 99 |
| 28135at9443 | Nodal growth differentiation factor |
| 2666at9443 | Synemin |
| 37831at9443 | Acrosomal vesicle protein 1 |
| 2988at9443 | ATP-dependent RNA helicase DHX29 |
| 44834at9443 | Immediate early response 5 |
| 47668at9443 | Charged multivesicular body protein 6 |
| 30038at9443 | Stromal cell derived factor 4 |
| 25457at9443 | Cathepsin E |
| 27862at9443 | Golgi reassembly stacking protein 1 |
| 30950at9443 | cAMP responsive element binding protein 3 |
| 50806at9443 | Coiled-coil-helix-coiled-coil-helix domain containing 7 |
| 44769at9443 | Methionine sulfoxide reductase B2 |
| 43029at9443 | RAB12, member RAS oncogene family |
| 18462at9443 | Lengsin, lens protein with glutamine synthetase domain |
| 45390at9443 | Centrin 1 |
| 39986at9443 | Membrane spanning 4-domains A15 |
| 40550at9443 | UBX domain protein 8 |
| 43326at9443 | High mobility group box 4 |
| 23685at9443 | Mitogen-activated protein kinase kinase 7 |
| 43427at9443 | Neuromedin S |
| 5280at9443 | Sodium/potassium-transporting ATPase subunit alpha |
| 45120at9443 | Killer cell lectin like receptor G1 |
| 21806at9443 | Fibrinogen gamma chain |
| 24347at9443 | Reticulon 4 interacting protein 1 |
| 45529at9443 | homeobox protein SEBOX |
| 36403at9443 | translational activator of cytochrome c oxidase 1 |
| 31508at9443 | olfactory receptor 52B2 |
| 50338at9443 | Keratin associated protein 7-1 (gene/pseudogene) |
| 17073at9443 | interferon regulatory factor 2-binding protein-like |
| 38648at9443 | Potassium channel regulator |
| 43434at9443 | ADP ribosylation factor like GTPase 4D |
| 22535at9443 | Family with sequence similarity 217 member A |
| 32769at9443 | diacylglycerol O-acyltransferase 2-like protein 6 |
| 38962at9443 | Interferon induced protein 35 |
| 43131at9443 | lymphocyte antigen 6 complex locus protein G5b |
| 25441at9443 | Solute carrier family 46 member 1 |
| 49259at9443 | testis-specific H1 histone |
| 14971at9443 | Delta like canonical Notch ligand 3 |
| 45472at9443 | CD70 antigen |
| 28868at9443 | GRINL1A complex locus 1 |
| 46409at9443 | NK3 homeobox 1 |
| 43634at9443 | Mitochondrial ribosomal protein L13 |
| 49864at9443 | platelet basic protein |
| 34768at9443 | Zinc finger CCHC-type containing 3 |
| 15766at9443 | conserved oligomeric Golgi complex subunit 8 |
| 31852at9443 | RNA polymerase III subunit D |
| 24373at9443 | Pleckstrin homology domain containing O2 |
| 33658at9443 | Relaxin/insulin like family peptide receptor 4 |
| 49199at9443 | Protein phosphatase 1 regulatory inhibitor subunit 14A |
| 50384at9443 | LOW QUALITY PROTEIN: prostate and breast cancer overexpressed gene 1 protein |
| 33143at9443 | taste receptor type 2 member 40 |
| 20784at9443 | Zinc finger protein 641 |
| 28344at9443 | Kelch domain containing 9 |
| 49091at9443 | Neuromedin U |
| 38639at9443 | Homeobox protein |
| 20937at9443 | Solute carrier family 2 member 2 |
| 32236at9443 | Starch binding domain 1 |
| 18944at9443 | Angiopoietin 1 |
| 17063at9443 | Beta-hexosaminidase |
| 33356at9443 | olfactory receptor 51A7 |
| 42547at9443 | Chromosome 18 open reading frame 21 |
| 10681at9443 | MYB proto-oncogene, transcription factor |
| 8373at9443 | Complement C7 |
| 11346at9443 | tubulin polyglutamylase TTLL11 |
| 24710at9443 | StAR related lipid transfer domain containing 3 |
| 46560at9443 | keratin-associated protein 11-1 |
| 7237at9443 | Kinesin family member 17 |
| 40104at9443 | NECAP endocytosis associated 1 |
| 22644at9443 | Aldehyde dehydrogenase 5 family member A1 |
| 23104at9443 | Ankyrin repeat and SOCS box containing 10 |
| 22303at9443 | sedoheptulokinase |
| 24487at9443 | bone morphogenetic protein 10 |
| 20120at9443 | RAN binding protein 3 like |
| 29432at9443 | Zinc activated ion channel |
| 44659at9443 | interferon beta |
| 30760at9443 | Phospholipid phosphatase related 5 |
| 50934at9443 | TP53-regulated inhibitor of apoptosis 1 |
| 50042at9443 | mediator of RNA polymerase II transcription subunit 9 |
| 38750at9443 | 2-oxoglutarate and iron dependent oxygenase domain containing 3 |
| 47146at9443 | Chromosome 9 open reading frame 116 |
| 21890at9443 | Cytochrome P450 family 27 subfamily B member 1 |
| 29946at9443 | Exo/endonuclease G |
| 49499at9443 | Chromosome 11 open reading frame 86 |
| 51498at9443 | potassium voltage-gated channel subfamily E regulatory beta subunit 5 |
| 35601at9443 | Aquaporin 5 |
| 40867at9443 | Phosphatidylcholine transfer protein |
| 46769at9443 | Mitochondrial ribosomal protein L17 |
| 6508at9443 | Smith-Magenis syndrome chromosome region, candidate 8 |
| 19484at9443 | Coiled-coil domain containing 183 |
| 31766at9443 | Decapping enzyme, scavenger |
| 28320at9443 | Cyclin J like |
| 25658at9443 | Sad1 and UNC84 domain containing 5 |
| 24432at9443 | Vasopressin V1a receptor |
| 40182at9443 | Leucine rich repeat containing 10B |
| 16693at9443 | Protein phosphatase 1 regulatory subunit 3F |
| 44513at9443 | protein TNT |
| 40013at9443 | protein MAK16 homolog |
| 37783at9443 | Chromosome 5 open reading frame 51 |
| 28351at9443 | Transmembrane protein 171 |
| 37030at9443 | Dolichyl-phosphate mannosyltransferase subunit 1, catalytic |
| 12859at9443 | Component of oligomeric golgi complex 6 |
| 44284at9443 | Neuronal calcium sensor 1 |
| 43377at9443 | Ferredoxin 2 |
| 38337at9443 | Proline rich 22 |
| 49292at9443 | Leucine rich single-pass membrane protein 2 |
| 12196at9443 | Electron transfer flavoprotein dehydrogenase |
| 45489at9443 | Paired like homeobox 2b |
| 25111at9443 | disrupted in renal carcinoma protein 2 |
| 25116at9443 | nuclear fragile X mental retardation-interacting protein 1 |
| 46150at9443 | Tetratricopeptide repeat domain 32 |
| 23522at9443 | carboxypeptidase E |
| 35703at9443 | Isochorismatase domain containing 1 |
| 40567at9443 | RAB11B, member RAS oncogene family |
| 30192at9443 | PNMA family member 1 |
| 30907at9443 | Annexin |
| 49072at9443 | Mediator of RNA polymerase II transcription subunit 11 |
| 9915at9443 | N-ethylmaleimide sensitive factor, vesicle fusing ATPase |
| 27725at9443 | phosphorylated adapter RNA export protein |
| 26687at9443 | Glucose-6-phosphatase |
| 40236at9443 | POU class 2 associating factor 1 |
| 38218at9443 | Lymphatic vessel endothelial hyaluronan receptor 1 |
| 19204at9443 | nuclear receptor subfamily 1 group D member 1 |
| 16430at9443 | REL proto-oncogene, NF-kB subunit |
| 19714at9443 | Coiled-coil domain containing 85A |
| 37129at9443 | Ubiquitin thioesterase |
| 37505at9443 | Integral membrane protein 2A |
| 50129at9443 | GADD45G interacting protein 1 |
| 43782at9443 | 5-formyltetrahydrofolate cyclo-ligase |
| 28213at9443 | Inhibin beta E subunit |
| 44218at9443 | Sodium voltage-gated channel beta subunit 4 |
| 45874at9443 | Titin-cap |
| 15122at9443 | striatin-4 isoform X1 |
| 43097at9443 | voltage-dependent calcium channel gamma-1 subunit |
| 15957at9443 | N-acetylglucosamine-6-sulfatase |
| 13338at9443 | rho GTPase-activating protein 40 |
| 41468at9443 | Myomaker, myoblast fusion factor |
| 19261at9443 | Zinc finger protein 219 |
| 892at9443 | Polycystin family receptor for egg jelly |
| 50274at9443 | Dynein light chain roadblock |
| 48702at9443 | Gametocyte specific factor 1 like |
| 39362at9443 | Hypoxanthine phosphoribosyltransferase |
| 14218at9443 | NDC80, kinetochore complex component |
| 45112at9443 | Achaete-scute family bHLH transcription factor 3 |
| 23002at9443 | keratin-associated protein 16-1 |
| 43960at9443 | Pre-mRNA processing factor 38B |
| 26847at9443 | Solute carrier family 46 member 2 |
| 38759at9443 | Family with sequence similarity 3 member B |
| 31713at9443 | keratin-associated protein 29-1 |
| 2993at9443 | ArfGAP with RhoGAP domain, ankyrin repeat and PH domain 1 |

| | |
|---|---|
| 38041at9443 | olfactory receptor 10K2 |
| 44543at9443 | Angiopoietin like 8 |
| 1784at9443 | DNA-directed RNA polymerase II subunit RPB1 |
| 22848at9443 | Phosphoserine aminotransferase |
| 43387at9443 | Sulfhydryl oxidase |
| 35878at9443 | Proteasome subunit beta type |
| 26689at9443 | SRY-box 7 |
| 48388at9443 | transmembrane protein 170B |
| 807at9443 | Voltage-dependent P/Q-type calcium channel subunit alpha |
| 20166at9443 | NADH dehydrogenase |
| 47911at9443 | otoraplin |
| 44836at9443 | Chromosome 17 open reading frame 98 |
| 41252at9443 | GrpE protein homolog |
| 16695at9443 | Family with sequence similarity 83 member C |
| 22358at9443 | Adaptor related protein complex 2 mu 1 subunit |
| 25065at9443 | fibrous sheath CABYR-binding protein |
| 7578at9443 | General transcription factor IIIC subunit 3 |
| 22612at9443 | zinc finger protein 597 |
| 42227at9443 | CD320 antigen isoform X1 |
| 33092at9443 | olfactory receptor 51S1 |
| 9380at9443 | threonine synthase-like 1 |
| 30508at9443 | Potassium two pore domain channel subfamily K member 18 |
| 48063at9443 | Caveolae associated protein 3 |
| 22001at9443 | Cilia and flagella associated protein 53 |
| 25562at9443 | N-acetylglucosamine-6-phosphate deacetylase |
| 10896at9443 | Zinc finger SWIM-type containing 3 |
| 16310at9443 | Werner helicase interacting protein 1 |
| 32984at9443 | Visual system homeobox 2 |
| 49736at9443 | histone H1t |
| 45761at9443 | translocator protein |
| 28608at9443 | Sperm associated antigen 8 |
| 42768at9443 | Biliverdin reductase B |
| 30349at9443 | POU domain, class 3, transcription factor 2 |
| 16319at9443 | RNA pseudouridylate synthase domain containing 2 |
| 42828at9443 | Guanylate cyclase activator 1C |
| 49861at9443 | Chromosome 20 open reading frame 202 |
| 29975at9443 | G protein-coupled receptor 139 |
| 25967at9443 | bone morphogenetic protein 15 |
| 30771at9443 | Homeobox C10 |
| 45405at9443 | zinc finger matrin-type protein 2 |
| 35982at9443 | TWIST neighbor |
| 25923at9443 | cornulin |
| 42901at9443 | Phosphomevalonate kinase |
| 50208at9443 | CART prepropeptide |
| 26831at9443 | Olfactomedin like 3 |
| 36934at9443 | Enoyl-CoA hydratase domain containing 2 |
| 48609at9443 | Chromosome 16 open reading frame 91 |
| 31194at9443 | retinol dehydrogenase 10 |
| 46179at9443 | Signaling threshold regulating transmembrane adaptor 1 |
| 22751at9443 | chitobiosyldiphosphodolichol beta-mannosyltransferase |
| 39869at9443 | L-xylulose reductase |
| 30420at9443 | SH3 and cysteine rich domain 2 |
| 48041at9443 | GON7, KEOPS complex subunit |
| 47116at9443 | Proprotein convertase subtilisin/kexin type 1 inhibitor |
| 14601at9443 | MAGE family member L2 |
| 37905at9443 | Insulin like growth factor binding protein 5 |
| 48714at9443 | mitochondrial fission 1 protein |
| 31960at9443 | Calcium homeostasis modulator family member 5 |
| 13534at9443 | Transporter |
| 17393at9443 | Basal cell adhesion molecule (Lutheran blood group) |
| 27222at9443 | growth/differentiation factor 8 |
| 24608at9443 | Protein O-glucosyltransferase 1 |
| 25961at9443 | Transmembrane protein with EGF like and two follistatin like domains 2 |
| 43139at9443 | Tetraspanin 13 |
| 44673at9443 | Transmembrane protein 52 |
| 27637at9443 | Protein phosphatase, Mg2+/Mn2+ dependent 1L |
| 41608at9443 | tumor necrosis factor ligand superfamily member 9 |
| 38514at9443 | Gamma-glutamylcyclotransferase |
| 49280at9443 | Prostate and testis expressed 1 |
| 42585at9443 | Cyclin dependent kinase 2 interacting protein |
| 36614at9443 | Non imprinted in Prader-Willi/Angelman syndrome 1 |
| 22769at9443 | Lipopolysaccharide binding protein |
| 31260at9443 | Caveolae associated protein 2 |
| 22833at9443 | AT-rich interaction domain 3B |
| 49759at9443 | Migration and invasion enhancer 1 |
| 5950at9443 | Aminopeptidase |
| 45924at9443 | BCL2 associated agonist of cell death |
| 19848at9443 | Calcitonin receptor |
| 48430at9443 | Serglycin |
| 9983at9443 | SPG7, paraplegin matrix AAA peptidase subunit |
| 46820at9443 | Achaete-scute family bHLH transcription factor 5 |
| 21881at9443 | tRNA methyltransferase 61B |
| 42152at9443 | glutathione peroxidase 2 |
| 33482at9443 | Solute carrier family 25 member 20 |
| 32575at9443 | Sorting nexin family member 21 |
| 25454at9443 | Protein Wnt |
| 10078at9443 | KIAA1522 |
| 49005at9443 | protein ripply3 |
| 17345at9443 | Breast cancer type 1 susceptibility protein homolog |
| 26907at9443 | Alcohol dehydrogenase 4 (class II), pi polypeptide |
| 42503at9443 | Distal-less homeobox 4 |
| 32026at9443 | junctional adhesion molecule 3 |
| 32835at9443 | Homeobox A5 |
| 21158at9443 | dolichol kinase |
| 29356at9443 | Zona pellucida binding protein 2 |
| 41108at9443 | UTP23, small subunit processome component |
| 30799at9443 | Solute carrier family 25 member 32 |
| 36086at9443 | interleukin-1 beta |
| 33409at9443 | hydroxycarboxylic acid receptor 1 |
| 23249at9443 | transmembrane protein 151A |
| 31223at9443 | Somatostatin receptor 4 |
| 4838at9443 | Nucleolar protein 6 |
| 44468at9443 | Fibroblast growth factor |
| 31393at9443 | Pentraxin 3 |
| 29684at9443 | Glutaminyl-peptide cyclotransferase |
| 34438at9443 | Cilia and flagella associated protein 77 |
| 23603at9443 | Fc receptor like 2 |
| 23569at9443 | DNA primase small subunit |
| 28628at9443 | Fructose-bisphosphate aldolase |
| 40639at9443 | Dual specificity phosphatase 19 |
| 39957at9443 | granzyme K |
| 39832at9443 | caspase-14 |
| 39554at9443 | AMMECR1 |
| 38621at9443 | out at first protein homolog |
| 6572at9443 | GRB2 associated regulator of MAPK1 subtype 1 |
| 44489at9443 | Thioredoxin domain containing 12 |
| 43137at9443 | Vitelline membrane outer layer 1 homolog |
| 50986at9443 | orexin |
| 29825at9443 | WD repeat domain 54 |
| 25811at9443 | Neuronal pentraxin 1 |
| 29620at9443 | Protein phosphatase 1 regulatory subunit 15A |
| 42549at9443 | noggin |
| 39896at9443 | Telomere repeat binding bouquet formation protein 2 |
| 34351at9443 | HAUS augmin-like complex subunit 7 |
| 29773at9443 | spermatogenic leucine zipper protein 1 |
| 37395at9443 | Prostaglandin E receptor 2 |
| 50487at9443 | C-X-C motif chemokine |
| 47105at9443 | MRG domain binding protein |
| 48788at9443 | nuclear transition protein 2 |
| 36159at9443 | myogenic differentiation 1 |
| 12936at9443 | Hermansky-Pudlak syndrome 6 protein |
| 33785at9443 | Zinc finger HIT-type containing 2 |
| 45736at9443 | Inhibitor of DNA binding 2 |
| 32896at9443 | Fos proto-oncogene, AP-1 transcription factor subunit |
| 51006at9443 | C-X-C motif chemokine ligand 17 |
| 6136at9443 | Catsper channel auxiliary subunit epsilon |
| 21035at9443 | Porcupine O-acyltransferase |
| 31347at9443 | Developing brain homeobox 2 |
| 33487at9443 | coiled-coil domain-containing protein 185 |
| 30663at9443 | XK related 8 |
| 26532at9443 | LOW QUALITY PROTEIN: DEP domain-containing protein 4 |
| 50689at9443 | Guanylate cyclase activator 2A |
| 51648at9443 | Fc fragment of IgE receptor Ig |
| 50703at9443 | COMM domain containing 6 |
| 36587at9443 | F-box protein 45 |
| 39130at9443 | Follistatin like 3 |
| 9372at9443 | ERCC excision repair 2, TFIIH core complex helicase subunit |
| 24891at9443 | Neuronal pentraxin 2 |
| 22847at9443 | Cholinergic receptor nicotinic alpha 1 subunit |
| 43560at9443 | Heat shock protein family B (small) member 8 |
| 37096at9443 | Limb and CNS expressed 1 |
| 44197at9443 | Protein lin-7 homolog |
| 24031at9443 | Chromosome 11 open reading frame 95 |
| 46503at9443 | Claudin |
| 42640at9443 | Transmembrane BAX inhibitor motif containing 4 |
| 34523at9443 | GLIPR1 like 2 |
| 20110at9443 | SAMM50 sorting and assembly machinery component |
| 41928at9443 | Fibroblast growth factor |
| 31034at9443 | IKBKB interacting protein |
| 42371at9443 | Heme binding protein 1 |
| 37440at9443 | Leucine rich repeat containing 73 |
| 27254at9443 | retrotransposon Gag like 3 |
| 34000at9443 | BCL2 associated athanogene 4 |
| 49371at9443 | Coiled-coil-helix-coiled-coil-helix domain containing 1 |
| 14303at9443 | Acyl-CoA dehydrogenase very long chain |
| 44991at9443 | Coiled-coil domain containing 184 |
| 47670at9443 | charged multivesicular body protein 4b |
| 39093at9443 | PGAM family member 5, mitochondrial serine/threonine protein phosphatase |
| 28719at9443 | zinc finger protein 296 |
| 23559at9443 | protein Red |
| 44208at9443 | Cerebellin 1 precursor |
| 46000at9443 | regulator of G-protein signaling 21 |
| 12690at9443 | Sirtuin 1 |
| 49217at9443 | somatostatin |
| 40558at9443 | Glycoprotein A33 |
| 48167at9443 | Chromosome 11 open reading frame 58 |
| 8444at9443 | Hepatocyte growth factor |
| 22040at9443 | Mitochondrial import inner membrane translocase subunit TIM44 |
| 24327at9443 | Tachykinin receptor 2 |
| 38609at9443 | Cytochrome b reductase 1 |
| 12036at9443 | Carnitine palmitoyltransferase 2 |
| 23394at9443 | Unc-51 like kinase 3 |
| 35215at9443 | Caveolae associated protein 1 |
| 19175at9443 | F-box only protein 33 |
| 40743at9443 | Chromosome 6 open reading frame 229 |
| 27103at9443 | probable G-protein coupled receptor 152 |
| 47380at9443 | Transmembrane protein 114 |
| 26966at9443 | 5-hydroxytryptamine receptor 1D |
| 13341at9443 | transcription initiation factor TFIID subunit 5 |
| 30524at9443 | Aquaporin 4 |
| 40672at9443 | TIMP metallopeptidase inhibitor 4 |
| 48201at9443 | natriuretic peptides A |
| 40107at9443 | Leucine rich repeat containing 38 |
| 28606at9443 | E2F transcription factor 2 |
| 19164at9443 | Aspartyl-tRNA synthetase |
| 2190at9443 | intron-binding protein aquarius |
| 33381at9443 | Family with sequence similarity 217 member B |
| 41995at9443 | rho GDP-dissociation inhibitor 3 |
| 45264at9443 | VPS37B, ESCRT-I subunit |
| 41933at9443 | Ephrin A1 |
| 47322at9443 | Cysteine rich hydrophobic domain 2 |
| 26709at9443 | Adrenoceptor alpha 2A |
| 7275at9443 | adhesion G-protein coupled receptor D2 |
| 48150at9443 | interleukin-9 |
| 29163at9443 | Alpha-1-microglobulin/bikunin precursor |
| 5423at9443 | Mtr4 exosome RNA helicase |
| 24084at9443 | Ubiquilin 1 |
| 39490at9443 | Family with sequence similarity 180 member B |
| 42003at9443 | Peroxisomal biogenesis factor 11 gamma |
| 38928at9443 | MKRN2 opposite strand protein |
| 43116at9443 | Potassium channel tetramerization domain containing 2 |
| 37680at9443 | BMP and activin membrane-bound inhibitor homolog |
| 26412at9443 | bystin |
| 37942at9443 | estradiol 17-beta-dehydrogenase 8 |
| 17177at9443 | nuclear receptor subfamily 1 group D member 2 |
| 24897at9443 | complement C1q tumor necrosis factor-related protein 3 isoform X1 |
| 43445at9443 | RAS related |
| 38510at9443 | fragile X mental retardation 1 neighbor protein |
| 32667at9443 | coiled-coil domain-containing protein 54 |
| 23545at9443 | Acetyl-CoA acetyltransferase 2 |
| 31529at9443 | Geminin coiled-coil domain containing |
| 37780at9443 | Zinc finger protein 688 |
| 39857at9443 | collagen alpha-1(XXIII) chain |
| 21645at9443 | lamin-B1 |
| 44035at9443 | NADH |
| 38186at9443 | Visual system homeobox 1 |
| 30107at9443 | torsin-1A |
| 47192at9443 | Interleukin 17A |

| | |
|---|---|
| 48659at9443 | EP300-interacting inhibitor of differentiation 2 |
| 31645at9443 | E2F transcription factor 4 |
| 32598at9443 | Leucine rich repeat containing 52 |
| 26873at9443 | Cartilage associated protein |
| 39678at9443 | Intercellular adhesion molecule 2 |
| 13013at9443 | retinitis pigmentosa 1-like 1 protein |
| 33879at9443 | protein FAM228B |
| 29068at9443 | Acrosin |
| 37388at9443 | Homeobox C13 |
| 24555at9443 | 5-hydroxytryptamine receptor 3B |
| 44835at9443 | BLOC-1 related complex subunit 6 |
| 48408at9443 | myotrophin |
| 34336at9443 | Biphenyl hydrolase like |
| 17803at9443 | Heparanase |
| 36872at9443 | deoxycytidine kinase |
| 30156at9443 | Caveolae associated protein 4 |
| 43903at9443 | Peptidyl-prolyl cis-trans isomerase |
| 22509at9443 | growth hormone-releasing hormone receptor |
| 19187at9443 | carboxylesterase 4A |
| 29047at9443 | G-protein coupled receptor 15 |
| 18053at9443 | Alkaline phosphatase |
| 25476at9443 | diphthamide biosynthesis 1 |
| 29486at9443 | dentin sialophosphoprotein |
| 46267at9443 | Cornichon family AMPA receptor auxiliary protein 2 |
| 38243at9443 | phenylethanolamine N-methyltransferase |
| 28474at9443 | Aminocarboxymuconate semialdehyde decarboxylase |
| 26176at9443 | Coagulation factor X |
| 45586at9443 | phospholipase A2 group IIE |
| 34526at9443 | Rab interacting lysosomal protein |
| 13849at9443 | Syntaxin binding protein 3 |
| 30495at9443 | Angiopoietin like 7 |
| 35918at9443 | olfactory receptor 52W1 |
| 38333at9443 | ELL-associated factor 1 |
| 30814at9443 | Trace amine associated receptor 5 |
| 42624at9443 | SPRY domain containing 4 |
| 28637at9443 | gastrin-releasing peptide receptor |
| 24106at9443 | forkhead box protein C2 |
| 38640at9443 | Mitochondrial ribosomal protein L16 |
| 37907at9443 | Centromere protein R |
| 28202at9443 | TNFAIP3 interacting protein 2 |
| 21438at9443 | carboxypeptidase A6 |
| 44880at9443 | Apolipoprotein D |
| 2777at9443 | maestro heat-like repeat family member 5 |
| 36039at9443 | Proteasome assembly chaperone 1 |
| 50200at9443 | Transmembrane protein 233 |
| 45302at9443 | Peptidyl-prolyl cis-trans isomerase |
| 33708at9443 | Ephrin B1 |
| 15696at9443 | tRNA wybutosine-synthesizing protein 4 |
| 36607at9443 | Mitochondrial ribosomal protein L46 |
| 42958at9443 | C-type lectin domain family 3 member B |
| 16622at9443 | Ankyrin repeat and ubiquitin domain containing 1 |
| 33612at9443 | Testis expressed 26 |
| 36652at9443 | forkhead box protein B1 |
| 41690at9443 | Homeobox B5 |
| 15895at9443 | catalase |
| 28239at9443 | Transcobalamin 1 |
| 1196at9443 | CREB binding protein |
| 36333at9443 | Dehydrogenase/reductase 13 |
| 36680at9443 | DNA-directed RNA polymerase II subunit RPB3 |
| 25782at9443 | ankyrin repeat and SOCS box protein 18 |
| 48770at9443 | N-acetyltransferase domain containing 1 |
| 43787at9443 | Claudin |
| 26844at9443 | Tachykinin receptor 3 |
| 25503at9443 | Arrestin domain containing 4 |
| 35252at9443 | homeobox protein aristaless-like 3 |
| 11453at9443 | acyl-CoA oxidase 1 |
| 14822at9443 | Oncoprotein induced transcript 3 |
| 40324at9443 | Forkhead box A3 |
| 25027at9443 | Cell cycle checkpoint control protein |
| 24679at9443 | Potassium voltage-gated channel subfamily A member 6 |
| 38919at9443 | taste receptor type 2 member 16 |
| 19158at9443 | Patatin like phospholipase domain containing 3 |
| 39504at9443 | Neurexophilin |
| 31447at9443 | G protein-coupled receptor 137C |
| 36769at9443 | NADH |
| 37487at9443 | SLAM family member 6 |
| 42792at9443 | transmembrane protein 247 |
| 45251at9443 | Transmembrane 4 L six family member 4 |
| 27145at9443 | DDB1- and CUL4-associated factor 7 |
| 5154at9443 | nestin |
| 26345at9443 | chloride intracellular channel protein 6 |
| 31879at9443 | ADP-ribosylhydrolase like 2 |
| 24415at9443 | Family with sequence similarity 71 member B |
| 38325at9443 | Asialoglycoprotein receptor 1 |
| 23292at9443 | WD repeat and SOCS box containing 2 |
| 48099at9443 | Mitochondrial ribosomal protein S14 |
| 43257at9443 | homeobox protein engrailed-2 |
| 23219at9443 | PHD finger protein 21B |
| 49223at9443 | Prokineticin 2 |
| 28858at9443 | Queuosine salvage protein |
| 41794at9443 | Phospholipase A2 group XIIB |
| 43100at9443 | Nucleoredoxin like 1 |
| 41659at9443 | mas-related G-protein coupled receptor member G |
| 28079at9443 | ethanolaminephosphotransferase 1 |
| 44773at9443 | neurogenin-1 |
| 30916at9443 | Forkhead box I1 |
| 27754at9443 | carbohydrate sulfotransferase 14 |
| 38561at9443 | Homeobox B8 |
| 11015at9443 | Actin filament associated protein 1 like 1 |
| 39576at9443 | Chromosome 3 open reading frame 70 |
| 35586at9443 | Neuropeptides B and W receptor 1 |
| 37439at9443 | Olfactory receptor |
| 38796at9443 | osteopetrosis-associated transmembrane protein 1 |
| 36814at9443 | carboxymethylenebutenolidase homolog |
| 19283at9443 | Muscarinic acetylcholine receptor |
| 49816at9443 | Succinate dehydrogenase complex assembly factor 4 |
| 32887at9443 | Leucine rich alpha-2-glycoprotein 1 |
| 7882at9443 | integrator complex subunit 5 |
| 36743at9443 | serine protease 27 |
| 38129at9443 | Transmembrane protein 38A |
| 29179at9443 | ceramide synthase 5 isoform X1 |
| 37074at9443 | Ankyrin repeat domain 60 |
| 31514at9443 | Cathepsin S |
| 40102at9443 | Kallikrein related peptidase 4 |

| | |
|---|---|
| 3426at9443 | Fibulin 2 |
| 39962at9443 | Barttin CLCNK type accessory beta subunit |
| 23600at9443 | Coiled-coil domain containing 105 |
| 31941at9443 | TNFAIP3-interacting protein 3 |
| 37072at9443 | Coiled-coil domain containing 189 |
| 38833at9443 | Transmembrane protein 53 |
| 22385at9443 | TNF receptor superfamily member 11b |
| 12383at9443 | synaptonemal complex protein 2-like |
| 42458at9443 | Homeobox C4 |
| 38973at9443 | Developing brain homeobox 1 |
| 28588at9443 | Armadillo repeat containing 12 |
| 12071at9443 | Arachidonate 15-lipoxygenase, type B |
| 35007at9443 | Sideroflexin |
| 40156at9443 | Nudix hydrolase 8 |
| 42202at9443 | RAB22A, member RAS oncogene family |
| 37825at9443 | TatD DNase domain containing 3 |
| 19192at9443 | prostacyclin synthase |
| 29483at9443 | Cholecystokinin B receptor |
| 40879at9443 | Receptor transporter protein 3 |
| 45038at9443 | Transmembrane 4 L six family member 5 |
| 33963at9443 | testis, prostate and placenta expressed |
| 32311at9443 | coiled-coil domain-containing protein 96 |
| 36346at9443 | Cerberus 1, DAN family BMP antagonist |
| 50815at9443 | C-C motif chemokine |
| 20770at9443 | Cytochrome P450 family 27 subfamily A member 1 |
| 32912at9443 | GPN-loop GTPase 2 |
| 40872at9443 | glutathione peroxidase 3 |
| 49029at9443 | Shisa like 2B |
| 9072at9443 | Chromosome 9 open reading frame 131 |
| 15615at9443 | Oxidative stress induced growth inhibitor family member 2 |
| 15970at9443 | Methyltransferase like 3 |
| 48301at9443 | Protein phosphatase 1 regulatory subunit 27 |
| 47507at9443 | endothelial cell-specific chemotaxis regulator |
| 7923at9443 | DNA replication licensing factor MCM6 |
| 19606at9443 | Acrosin binding protein |
| 28722at9443 | peroxisomal biogenesis factor 7 |
| 36960at9443 | aquaporin-1 |
| 33208at9443 | Cyclin N-terminal domain containing 1 |
| 35781at9443 | Olfactory receptor |
| 51464at9443 | tumor necrosis factor receptor superfamily member 13C |
| 17525at9443 | DNA-(apurinic or apyrimidinic site) lyase |
| 14214at9443 | Kelch like family member 21 |
| 18473at9443 | Matrix metallopeptidase 21 |
| 42321at9443 | Inhibitor of growth protein |
| 43933at9443 | PRA1 family protein |
| 45445at9443 | putative claudin-25 |
| 48757at9443 | Complexin 4 |
| 38137at9443 | Homeobox D1 |
| 9311at9443 | NOP2/Sun RNA methyltransferase family member 2 |
| 6465at9443 | DNA mismatch repair protein |
| 27980at9443 | Cytidine/uridine monophosphate kinase 2 |
| 44721at9443 | Myosin light chain, phosphorylatable, fast skeletal muscle |
| 1897at9443 | ATP binding cassette subfamily A member 3 |
| 45156at9443 | Cerebellin 4 precursor |
| 26456at9443 | Transmembrane 6 superfamily member 2 |
| 42417at9443 | Pancreatic and duodenal homeobox 1 |
| 18662at9443 | Tigger transposable element derived 5 |
| 21633at9443 | UBX domain protein 11 |
| 41538at9443 | keratin-associated protein 27-1 |
| 10023at9443 | Cyclin F |
| 40749at9443 | Pleckstrin homology domain containing B1 |
| 48346at9443 | apolipoprotein C-IV |
| 35928at9443 | C-X-C motif chemokine receptor 3 |
| 41575at9443 | testis expressed 35 |
| 28119at9443 | Four jointed box 1 |
| 50948at9443 | Regulator of cell cycle |
| 26351at9443 | calreticulin-3 |
| 44080at9443 | Glial cell derived neurotrophic factor |
| 6637at9443 | DDHD domain containing 1 |
| 26162at9443 | Receptor for activated C kinase 1 |
| 35300at9443 | Hydroxysteroid 17-beta dehydrogenase 3 |
| 13764at9443 | Collagen type IV alpha 1 chain |
| 25631at9443 | Mex-3 RNA binding family member B |
| 39033at9443 | homeobox protein orthopedia |
| 39241at9443 | TIMP metallopeptidase inhibitor 3 |
| 15395at9443 | ankyrin repeat and death domain containing 1A |
| 51220at9443 | protein S100-P |
| 27374at9443 | WD repeat domain 73 |
| 20642at9443 | Ubiquinol-cytochrome c reductase core protein 1 |
| 49763at9443 | Glutaredoxin 5 |
| 25524at9443 | Tryptophan 2,3-dioxygenase |
| 34783at9443 | Aspartate dehydrogenase domain containing |
| 16791at9443 | Acid sensing ion channel subunit family member 4 |
| 48919at9443 | Urotensin 2 |
| 6294at9443 | melanoma-associated antigen E1 |
| 38702at9443 | taste receptor type 2 member 4 |
| 19451at9443 | Drebrin 1 |
| 22095at9443 | APC down-regulated 1 like |
| 46904at9443 | Regulator of G protein signaling 9 binding protein |
| 37086at9443 | voltage-dependent calcium channel gamma-8 subunit |
| 39616at9443 | Homeobox B2 |
| 38016at9443 | Transmembrane protein 64 |
| 46212at9443 | neuroglobin |
| 24946at9443 | Acid phosphatase, prostate |
| 7810at9443 | Fc receptor like 5 |
| 43232at9443 | Crystallin beta A2 |
| 39902at9443 | Fibroblast growth factor |
| 46520at9443 | heat shock protein beta-3 |
| 34276at9443 | Insulin like growth factor binding protein 2 |
| 36243at9443 | Proline rich 32 |
| 40521at9443 | splicing factor 3B subunit 4 |
| 18907at9443 | platelet glycoprotein V |
| 41763at9443 | Peptidyl-prolyl cis-trans isomerase |
| 27245at9443 | exosome complex component RRP45 |
| 44022at9443 | ventral anterior homeobox 2 |
| 41679at9443 | Sjogren syndrome/scleroderma autoantigen 1 |
| 47619at9443 | Fer3 like bHLH transcription factor |
| 10074at9443 | Arachidonate 12-lipoxygenase, 12R type |
| 42683at9443 | Protein tyrosine phosphatase receptor type C-associated protein |
| 25900at9443 | SET and MYND domain containing 2 |
| 24916at9443 | Fc receptor like 1 |
| 24850at9443 | meiotic recombination protein SPO11 isoform X1 |
| 19543at9443 | nuclear receptor subfamily 0 group B member 1 |
| 14094at9443 | DBF4 zinc finger |

| | |
|---|---|
| 37017at9443 | dopamine receptor D4 |
| 39839at9443 | Spermatogenesis associated 9 |
| 25929at9443 | Centromere protein T |
| 44750at9443 | chromosome 17 open reading frame 107 |
| 45686at9443 | transmembrane protein 47 |
| 47189at9443 | Endonuclease G |
| 47536at9443 | Troponin C2, fast skeletal type |
| 25213at9443 | Cell cycle control protein |
| 13799at9443 | Amyloid beta precursor like protein 2 |
| 12867at9443 | Extracellular matrix protein 2 |
| 18237at9443 | Protein disulfide isomerase family A member 5 |
| 29428at9443 | Pannexin |
| 12330at9443 | Polypeptide N-acetylgalactosaminyltransferase |
| 25931at9443 | Abhydrolase domain containing 12B |
| 26418at9443 | GRINL1A complex locus 1 |
| 44858at9443 | Kruppel like factor 14 |
| 38361at9443 | TATA-box binding protein associated factor 9b |
| 15844at9443 | Tyrosine-protein kinase |
| 4099at9443 | Ceruloplasmin |
| 20973at9443 | Essential meiotic structure-specific endonuclease 1 |
| 50229at9443 | COX19, cytochrome c oxidase assembly factor |
| 17829at9443 | Mitochondrial elongation factor 2 |
| 33977at9443 | Ribonuclease H2 subunit B |
| 39559at9443 | glutathione peroxidase 6 |
| 26827at9443 | Sorting nexin |
| 45243at9443 | protein jagunal homolog 1 |
| 27098at9443 | Adrenoceptor alpha 2B |
| 39292at9443 | Rho family GTPase 1 |
| 46711at9443 | Epithelial mitogen |
| 50850at9443 | Galanin and GMAP prepropeptide |
| 34927at9443 | Potassium channel tetramerization domain containing 12 |
| 9249at9443 | AP-1 complex subunit gamma |
| 32377at9443 | Elongation factor Ts, mitochondrial |
| 17220at9443 | Activating transcription factor 6 beta |
| 42698at9443 | Shisa like 2A |
| 16409at9443 | Leucine rich repeats and calponin homology domain containing 4 |
| 26075at9443 | Abhydrolase domain containing 15 |
| 42039at9443 | N-6 adenine-specific DNA methyltransferase 1 |
| 20855at9443 | Ectonucleoside triphosphate diphosphohydrolase 6 (putative) |
| 27409at9443 | Protein Wnt |
| 40903at9443 | Tissue factor |
| 29482at9443 | Plasmalemma vesicle associated protein |
| 14239at9443 | dopamine beta-hydroxylase |
| 19835at9443 | negative elongation factor B |
| 38608at9443 | Radial spoke head 9 homolog |
| 31476at9443 | TEA domain transcription factor 4 |
| 37967at9443 | transmembrane emp24 domain-containing protein 6 |
| 43938at9443 | Ladybird homeobox 2 |
| 32569at9443 | S-methyl-5'-thioadenosine phosphorylase |
| 23735at9443 | CCR4-NOT transcription complex subunit 11 |
| 10820at9443 | Sulfhydryl oxidase |
| 8709at9443 | aryl hydrocarbon receptor |
| 29677at9443 | Paraoxonase 3 |
| 21101at9443 | tRNA wybutosine-synthesizing protein 2 homolog |
| 29843at9443 | protein FAM117B |
| 29153at9443 | Elongation of very long chain fatty acids protein 2 |
| 23970at9443 | Glucagon receptor |
| 31003at9443 | OTU deubiquitinase 1 |
| 31937at9443 | C-X-C motif chemokine receptor 5 |
| 43597at9443 | Interleukin 17C |
| 10518at9443 | ArfGAP with coiled-coil, ankyrin repeat and PH domains 3 |
| 19163at9443 | zinc finger protein 689 |
| 27307at9443 | Coagulation factor II thrombin receptor like 2 |
| 45516at9443 | WASH complex subunit 3 |
| 41155at9443 | Cysteine dioxygenase type 1 |
| 24531at9443 | C-type lectin domain containing 14A |
| 12528at9443 | CD248 molecule |
| 17432at9443 | G protein-coupled receptor 107 |
| 10424at9443 | Acyloxyacyl hydrolase |
| 35245at9443 | olfactory receptor 51G2 |
| 45348at9443 | Mitochondrial ribosomal protein S25 |
| 21823at9443 | lectin, mannose binding 1 like |
| 50907at9443 | 28 kDa heat- and acid-stable phosphoprotein |
| 28004at9443 | ST6 N-acetylgalactosaminide alpha-2,6-sialyltransferase 2 |
| 41363at9443 | Anti-silencing function 1A histone chaperone |
| 27320at9443 | Bombesin receptor subtype 3 |
| 43952at9443 | Insulin like growth factor binding protein 6 |
| 33350at9443 | Mitochondrial glycine transporter |
| 51164at9443 | C-C motif chemokine |
| 28479at9443 | Cell cycle control protein |
| 42149at9443 | Gap junction protein |
| 30239at9443 | Talin rod domain containing 1 |
| 35891at9443 | chymotrypsin-like elastase family member 1 |
| 6474at9443 | THAP domain containing 9 |
| 23940at9443 | SH2 domain-containing adapter protein B |
| 42790at9443 | myogenic factor 6 |
| 38042at9443 | zinc finger protein 784 |
| 26383at9443 | Galactose-1-phosphate uridylyltransferase |
| 50642at9443 | GTP cyclohydrolase I feedback regulator |
| 43494at9443 | mitochondrial assembly of ribosomal large subunit protein 1 |
| 28983at9443 | Layilin |
| 28765at9443 | MLLT3, super elongation complex subunit |
| 10509at9443 | TSR1, ribosome maturation factor |
| 24232at9443 | Acid phosphatase 2, lysosomal |
| 10788at9443 | Spermatogenesis associated 5 like 1 |
| 6093at9443 | Spalt like transcription factor 4 |
| 23252at9443 | Serum amyloid A like 1 |
| 39072at9443 | JunB proto-oncogene, AP-1 transcription factor subunit |
| 3435at9443 | Valosin containing protein interacting protein 1 |
| 18213at9443 | G protein-coupled receptor kinase |
| 27011at9443 | GDNF family receptor alpha-3 |
| 26033at9443 | Four and a half LIM domains 1 |
| 43565at9443 | transmembrane protein 223 |
| 29797at9443 | Protein phosphatase 1 regulatory subunit |
| 34562at9443 | keratin 9 |
| 51454at9443 | small integral membrane protein 13 |
| 50715at9443 | Small leucine rich protein 1 |
| 45363at9443 | Chromosome 1 open reading frame 185 |
| 39752at9443 | Tyrosine 3-monooxygenase/tryptophan 5-monooxygenase activation protein gamma |
| 30428at9443 | E2F transcription factor 1 |
| 36758at9443 | Zinc finger FYVE-type containing 21 |
| 16164at9443 | Pleckstrin homology domain containing N1 |
| 2950at9443 | Carboxypeptidase D |
| 4362at9443 | Leucine rich repeats and immunoglobulin like domains 3 |
| 36211at9443 | Nicotinate-nucleotide pyrophosphorylase |
| 22721at9443 | major centromere autoantigen B |
| 43678at9443 | signal peptidase complex subunit 3 |
| 37185at9443 | Glutamate-cysteine ligase modifier subunit |
| 22943at9443 | Mannosyltransferase |
| 5354at9443 | RAD51-associated protein 2 |
| 26304at9443 | Transmembrane 7 superfamily member 2 |
| 28208at9443 | Transmembrane and coiled-coil domains 6 |
| 11069at9443 | Zinc finger protein 792 |
| 37169at9443 | probable threonine protease PRSS50 |
| 16806at9443 | Sorting nexin |
| 43679at9443 | Calmodulin like 6 |
| 37699at9443 | Homeobox A11 |
| 44609at9443 | Histidine triad nucleotide binding protein 3 |
| 24983at9443 | Keratin 74 |
| 44243at9443 | transmembrane and coiled-coil domain-containing protein 2 |
| 44736at9443 | peptidoglycan recognition protein 1 |
| 42340at9443 | Cyclin dependent kinase inhibitor 1B |
| 48103at9443 | Fatty acid binding protein 2 |
| 35078at9443 | olfactory receptor 10AD1 |
| 49583at9443 | DR1 associated protein 1 |
| 41474at9443 | Shisa family member 3 |
| 37860at9443 | Polycomb group ring finger 1 |
| 42506at9443 | BRICHOS domain containing 5 |
| 40263at9443 | SIX homeobox 6 |
| 26504at9443 | solute carrier family 49 member 3 |
| 47920at9443 | Transthyretin |
| 38731at9443 | Insulin like growth factor binding protein 4 |
| 43841at9443 | Brain specific homeobox |
| 40171at9443 | replication protein A 30 kDa subunit |
| 21363at9443 | Docking protein 1 |
| 11643at9443 | Ribosome biogenesis protein BOP1 |
| 7366at9443 | Nuclear factor of activated T cells 2 |
| 39851at9443 | SLAM family member 8 |
| 10128at9443 | Trichohyalin like 1 |
| 38029at9443 | BARX homeobox 2 |
| 18569at9443 | Aldehyde dehydrogenase 8 family member A1 |
| 22918at9443 | glucagon-like peptide 1 receptor |
| 29218at9443 | Protein Wnt |
| 36723at9443 | calbindin |
| 34725at9443 | LysM domain containing 3 |
| 22511at9443 | Zinc finger protein 514 |
| 32733at9443 | ankyrin repeat and SOCS box protein 17 |
| 43287at9443 | RING finger protein 186 |
| 34696at9443 | Beta-1,3-N-acetylglucosaminyltransferase |
| 25548at9443 | Cytochrome P450 family 2 subfamily S member 1 |
| 41184at9443 | RAS like family 11 member B |
| 9669at9443 | Tripartite motif containing 71 |
| 36135at9443 | MAD2L1 binding protein |
| 45965at9443 | dCTP pyrophosphatase 1 |
| 10614at9443 | Chloride channel protein |
| 18643at9443 | MAGE family member E2 |
| 19763at9443 | Matrix metallopeptidase 19 |
| 33752at9443 | NFKB inhibitor alpha |
| 40502at9443 | regulated endocrine-specific protein 18 |
| 23730at9443 | DnaJ heat shock protein family (Hsp40) member A3 |
| 10389at9443 | Death domain containing 1 |
| 27438at9443 | Podocalyxin |
| 38802at9443 | Nudix hydrolase 21 |
| 31986at9443 | Stanniocalcin 2 |
| 22776at9443 | HtrA serine peptidase 1 |
| 47434at9443 | RING finger protein 11 |
| 25950at9443 | Growth differentiation factor 6 |
| 32222at9443 | Gamma-glutamyl hydrolase |
| 26071at9443 | Arrestin domain containing |
